# Ultra-Long-Term Delivery of Hydrophilic Drugs Using Injectable *In Situ* Cross-Linked Depots

**DOI:** 10.1101/2023.11.04.565631

**Authors:** Sohyung Lee, Spencer Zhao, Weihua Jiang, Xinyang Chen, Lingyun Zhu, John Joseph, Eli Agus, Helna Baby Mary, Shumaim Barooj, Kai Slaughter, Krisco Cheung, James N Luo, Chetan Shukla, Jingjing Gao, Dongtak Lee, Biji Balakrishnan, Christopher Jiang, Amogh Gorantla, Sukyung Woo, Jeffrey M Karp, Nitin Joshi

## Abstract

Achieving ultra-long-term release of hydrophilic drugs over several months remains a significant challenge for existing long-acting injectables (LAIs). Existing platforms, such as *in situ* forming implants (ISFI), exhibit high burst release due to solvent efflux and microsphere-based approaches lead to rapid drug diffusion due to significant water exchange and large pores. Addressing these challenges, we have developed an injectable platform that, for the first time, achieves ultra-long-term release of hydrophilic drugs for over six months. This system employs a methacrylated ultra-low molecular weight pre-polymer (polycaprolactone) to create *in situ* cross-linked depots (ISCD). The ISCD’s solvent-free design and dense mesh network, both attributed to the ultra-low molecular weight of the pre-polymer, effectively minimizes burst release and water influx/efflux. *In vivo* studies in rats demonstrate that ISCD outperforms ISFI by achieving lower burst release and prolonged drug release. We demonstrated the versatility of ISCD by showcasing ultra-long-term delivery of several hydrophilic drugs, including antiretrovirals (tenofovir alafenamide, emtricitabine, abacavir, and lamivudine), antibiotics (vancomycin and amoxicillin) and an opioid antagonist naltrexone. Additionally, ISCD achieved ultra-long-term release of the hydrophobic drug tacrolimus and enabled co-delivery of hydrophilic drug combinations encapsulated in a single depot. We also identified design parameters to tailor the polymer network, tuning drug release kinetics and ISCD degradation. Pharmacokinetic modeling predicted over six months of drug release in humans, significantly surpassing the one-month standard achievable for hydrophilic drugs with existing LAIs. The platform’s biodegradability, retrievability, and biocompatibility further underscore its potential for improving treatment adherence in chronic conditions.

## Introduction

Patient adherence to medications is a major obstacle to achieving effective treatment of numerous diseases, especially for chronic conditions that require treatment throughout life.^1–3^ Long-acting injectables (LAIs) simplify dosing schedules and enhance treatment regimen adherence^4–11^, which is particularly advantageous in low-resource settings with limited healthcare infrastructure.^9, 12–14^ Multiple LAIs are in clinical use for the treatment and prevention of different diseases.^15, 16^ Conventional LAI approaches such as microparticles,^17^ *in situ*-forming implants (ISFI), ^12, 18^ and wet milled particles^6, 15, 19–21^ have been documented to achieve prolonged release of hydrophobic drugs for several months but have poor ability to achieve similar ultra-long-term release of hydrophilic drugs^18, 22–24^. In the case of ISFI, which typically consists of a hydrophobic polymer, poly(lactic-co-glycolic acid) (PLGA) dissolved in a solvent, the solvent efflux during phase conversion tends to release a significant amount of drug as initial burst, which increases with the hydrophilicity of the drug.^18^ Additionally, previously developed LAI approaches, including ISFI and microspheres promote significant influx and efflux of water due to their large pores, leading to fast diffusion of hydrophilic drugs.^12, 17, 18^ Wet milling – a commonly employed approach for formulating LAIs^19, 20^ requires the drug to be hydrophobic and are fundamentally incompatible with hydrophilic drugs.^6, 15, 21^ Although implantable devices have shown success for long-term delivery of both hydrophobic^25^ and hydrophilic drugs^9, 14^, they require invasive, time-consuming medical procedures for insertion, which may pose significant challenges, particularly in low resource settings and low-middle income countries. Furthermore, invasive placement of implants often lead to a higher incidence of local inflammation compared to injectable alternatives, which typically involve less tissue disruption.^26, 27^

Hydrophilic drugs, which we defined as those with a solubility greater than 0.1 mg/mL, constitute a major fraction of all medications used for the prevention, treatment, and management of chronic conditions. Examples include anti-psychotics, anti-depressants, anti-convulsants, antibiotics, and drugs for treating substance abuse disorder (SUD). Although a few hydrophilic drugs have clinically approved LAI formulations, their release typically lasts for only a month. For example, Vivitrol®, an LAI suspension of naltrexone-loaded PLGA microspheres for treating SUD, provides 30 days of drug release,^28^ which is sub-optimal since SUD often requires therapy for several years.^29^ Therefore, there is an unmet need to develop an injectable platform that enables ultra-long-term delivery of hydrophilic drugs for several months. Additionally, the platform should be designed to be biodegradable for safe breakdown and clearance from the body while also being retrievable in the event of local or systemic drug toxicity.

We report an injectable platform **(Figure 1)** that addresses the limitations of current LAI approaches, enabling ultra-long-term release of hydrophilic drugs for at least 6 months. This platform is derived from an ultra-low-molecular-weight liquid pre-polymer – polycaprolactone (PCL, 500 < M_n_ < 2000), which has been used in multiple FDA-approved products.{Malikmammadov, 2018 #152} PCL is chemically modified with methacrylate groups **(Figure S1A)**, resulting in PCL dimethacrylate (PCLDMA) or PCL trimethacrylate (PCTMA), which can undergo free-radical polymerization in the presence of an appropriate initiator and accelerator. The liquid pre-polymer can effectively suspend or dissolve both hydrophilic and hydrophobic drugs, and can be easily injected through a standard 18-23 gauge needle **(Figure 1A)**. Upon co-injection with a clinically used radical initiator and an accelerator, benzoyl peroxide (BPO) and N,N-dimethylparatoluidine (DMT),^30–32^ respectively, the pre-polymer mixture undergoes a time-dependent cross-linking. This process physically encapsulates the drugs, creating a solid structure hereafter referred to as an *in situ* cross-linked depot (ISCD). Hydrolysis of the polymer ester bonds allows gradual erosion of the depot over time, obviating the need for surgical removal after depot exhaustion, and ensuring safe clearance from the body **(Figure S2)**.

**Figure 1.**
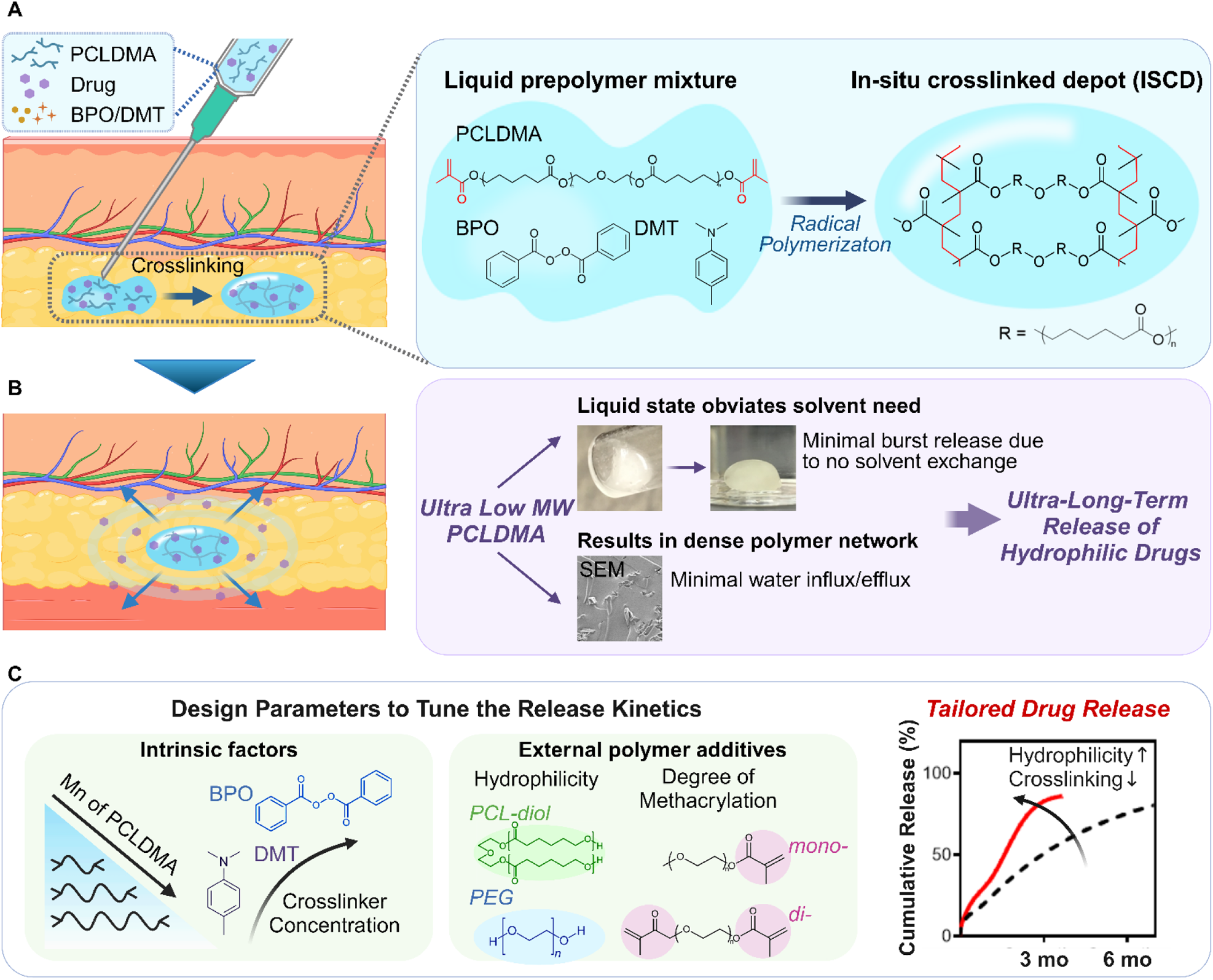
Injectable *in situ* crosslinked depot (ISCD) platform for sustained release of hydrophilic therapeutics. **A.** The main component of ISCD is low molecular weight liquid methacrylated PCL, for example, PCLDMA. The liquid pre-polymer can suspend or dissolve both hydrophilic and hydrophobic drugs and can be easily injected through a standard 18-23 gauge needle. Upon adding an initiator (BPO) and accelerator (DMT) to PCLDMA, the pre-polymer mixture undergoes radical polymerization transitioning from a free-flowing liquid solution to a solid monolithic depot, resulting in physical encapsulation of the drug. **B.** ISCD has two key features enabling ultra-long-term release of hydrophilic drugs: a solvent-free design and a dense mesh network, both attributed to the ultra-low-molecular weight of the pre-polymer, PCLDMA. The liquid state of the pre-polymer obviates the need for a solvent, minimizing burst release. Cross-linking of the ultra-small chains of the pre-polymer results in a dense network (as shown in the SEM image of an ISCD depot formed *in vitro*) that limits water influx and efflux, minimizing the drug release rate. **C.** Design parameters to tailor the ISCD network to tune the drug release kinetics. Modulating the intrinsic factors, including decreasing the concentrations of BPO and DMT, or using higher molecular weight PCLDMA increases drug release. Additionally, adding external polymer additives including polyethylene glycol (PEG) and PCL-diol alongside PCLDMA can enhance the depot’s hydrophilicity, increasing drug release. External additives with different degrees of methacrylation (mono or di) can further tune the drug release. The cumulative release profiles of TAF from two different ISCD depots injected subcutaneously into rats are shown as examples of tailored drug release with varying release rates. Lowering the crosslinking density and increasing the hydrophilicity of the polymer chains achieve faster drug release.

ISCD has two key features, which enable ultra-long-term release of hydrophilic drugs **(Figure 1B)**. These include a solvent-free design and a dense mesh network, both attributed to the use of ultra-low-molecular weight PCL. The liquid state of the pre-polymer eliminates the need for a solvent, minimizing the risk of high burst release, which is commonly associated with solvent exchange processes in ISFI.^24^ Additionally, the ultra-low molecular weight of methacrylated PCL forms a dense mesh upon cross-linking, which limits water influx/efflux, thereby controlling the drug release. We demonstrated that ISCD can exhibit sustained release of multiple hydrophilic drugs with varying water solubilities as high as 100 mg/mL and belonging to diverse therapeutic class, including antiretrovirals, opioid antagonists, and antibiotics. ISCD showed sustained release of all the drugs for at least 6-10 months *in vitro*, suggesting the versatility of the platform. We also demonstrated ultra-long-term release of two proof-of-concept hydrophilic drugs - tenofovir alafenamide (TAF) and naltrexone (NAL) - *in vivo* in rats. A single subcutaneous injection of ISCD formulations loaded with TAF or NAL resulted in sustained plasma concentration for at least 6-7 months. ISCD outperformed the conventional ISFI platform in TAF delivery, exhibiting notably lower burst release compared to ISFI, and providing a prolonged drug release duration of up to 7 months. ISCD also showed ultra-long-term release of a hydrophobic drug – Tacrolimus (TAC) - for at least 6 months in rats. Pharmacokinetic (PK) modeling predicted that the ISCD platform can achieve ultra-long-term delivery of naltrexone (NAL) and tacrolimus (TAC) for several months in humans. We also identified design parameters that can tailor the polymer network to tune the drug release kinetics and degradation of ISCD **(Figure 1C)**. Notably, modulating intrinsic factors, such as decreasing the concentrations of BPO and DMT or using a higher molecular weight of methacrylated PCL, increases drug release. Additionally, integrating external hydrophilic polymeric additives including polyethylene glycol (PEG) alongside methacrylated PCL can enhance drug release and depot degradation rate, which can be further fine-tuned by varying the degree of methacrylation (mono versus di) of the polymer additive. Finally, we also demonstrated the feasibility of achieving ultra-long-term release of clinically relevant combination regimens of hydrophilic drugs, encapsulated in a single depot, and demonstrated the biocompatibility and retrievability of ISCD.

Together, our findings underscore the potential of ISCD as a versatile platform to enable ultra-long-term delivery of both hydrophilic and hydrophobic drugs. To our knowledge, this is the first report demonstrating ultra-long-term delivery (>6 months) of hydrophilic drugs using an injectable system, significantly surpassing the standard of 1-month release avhievable by existing LAI approaches for hydrophilic drugs. This innovation holds the potential to revolutionize therapy options for a variety of chronic conditions where patient adherence is critical.

## Results

### Synthesis and characterization of ISCD

The main component of ISCD is ultra-low molecular weight methacrylated PCL (500 < M_n_ < 2000). PCL-diol or - triol were methacrylated *via* a reaction with methacrylic anhydride (MAA) and triethylamine (TEA), resulting in PCLDMA or PCLTMA, respectively **(Figure S1A)**. Methacrylation was confirmed via ^1^H NMR spectroscopy, with the ratio of integral areas under the protons of the double bond in the methacrylate group at δ = 6.12 ppm (2H, olefinic, cis) to the methylene protons of the PEG segment at δ = 3.70 ppm (4H, -OCH2CH2OCH2CH2O-) determined to be 1:2 **(Figure S1B)**.

Upon adding BPO and DMT to PCLDMA or PCLTMA, the pre-polymer mixture undergoes radical polymerization transitioning from a free-flowing liquid solution to a solid monolithic depot. The time interval between the mixing of different components and the complete crosslinking of ISCD is a critical parameter for successful clinical application. Ideally, the crosslinking kinetics should provide sufficient time to mix and inject the formulation before it solidifies. Therefore, we investigated the cross-linking time of ISCD, focusing on the impact of initiator and accelerator concentrations on it. Using a rotational rhemometer, we measured the cross-linking time as the point when the formulation’s viscosity starts to rapidly increase. When using 0.1 wt% each of BPO and DMT (BPO/DMT), the formulation demonstrated complete solidification within approximately 9 minutes **(Figure 2A)**. Increasing the concentrations of BPO/DMT from 0.1 to 0.3 wt% significantly reduced the PCLDMA crosslinking time to approximately 5 minutes, which further reduced to 2 minutes with BPO/DMT concentrations of 0.5 wt%. For subsequent experiments, we used 0.3 wt% concentrations of BMO/DMT.

**Figure 2.**
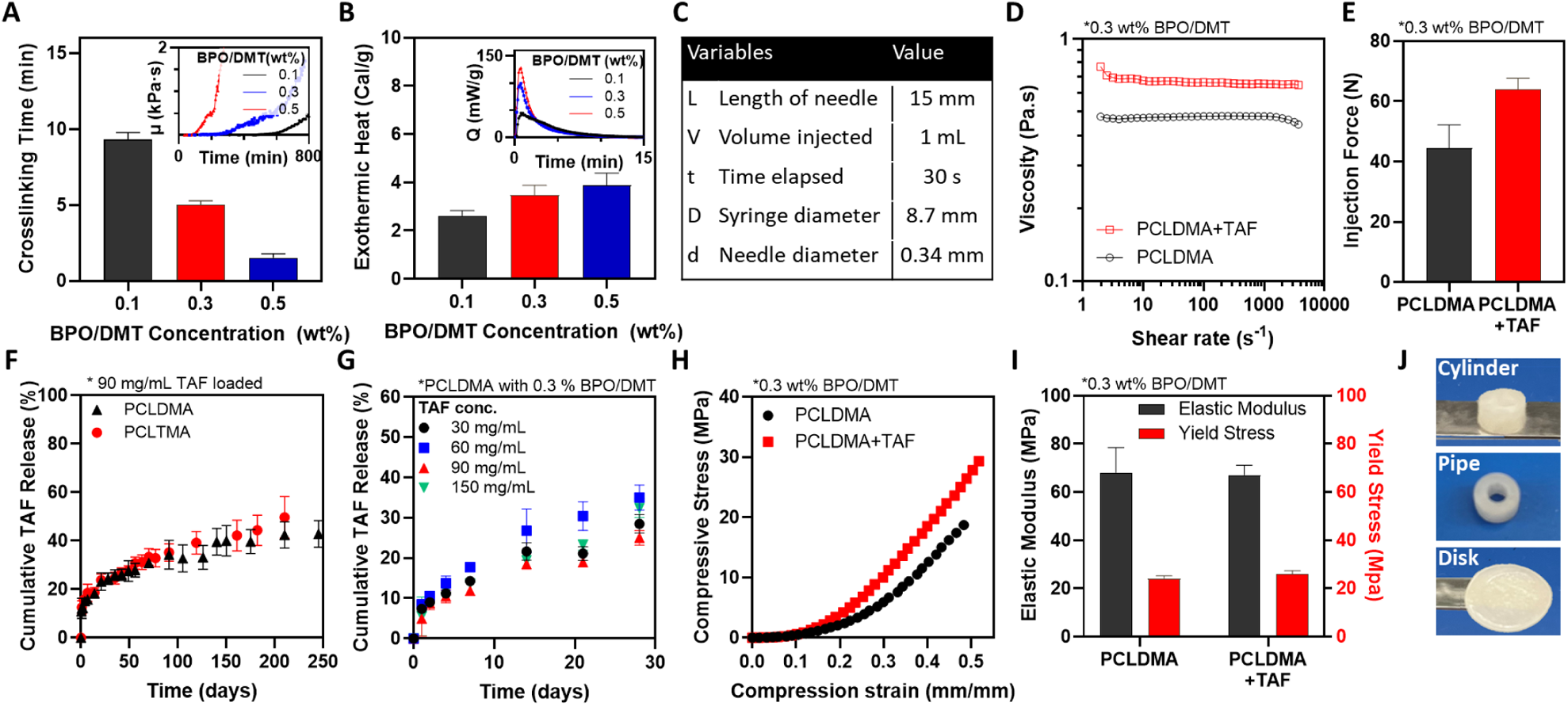
Synthesis and physiochemical characterization of the ISCD platform. **A.** The cross-linking time for PCLDMA at different BPO/DMT concentrations is shown. Cross-linking time was measured as the point at which the viscosity of the pre-polymer mixture, monitored with a rheometer, begins to increase rapidly, as shown in the inset. **B.** Exothermic heat released during the cross-linking of ISCD with varying concentrations of BPO/DMT, measured using DSC. **C.** Injection parameters for the Hagen-Poiseuille equation used to calculate the injection force for PCLDMA, with or without TAF, using a 23-gauge needle. **D.** Viscosities of PCLDMA, with or without TAF, measured using a rheometer. **E.** Injection force calculated for PCLDMA, with or without TAF. The maximum acceptable injection force is 80 N. **F.** *In vitro* release profile of TAF in PBS (37°C) from ISCD comprising either PCLDMA or PCLTMA. **G.** *In vitro* release profile of TAF in PBS (37°C) from ISCD loaded with concentrations of TAF. **H.** Compressive stress-strain curves for ISCD with or without TAF, measured using a mechanical tester. **I.** Elastic moduli and yield stress for ISCD with or without TAF. **J.** ISCD can be polymerized *ex vivo* into various shapes such as cylinders, pipes, or disks that can be used as ultra-long-acting implants. Data in A, B, E-G, and I are presented as mean ± standard deviation (n=3, replicates performed at least twice). Data in D and H are representative of a single experiment (repeated three times).

Since radical polymerization is an exothermic process, we intended to confirm whether *in situ* crosslinking of ISCD would result in any thermal tissue damage. We employed two complementary techniques: differential scanning calorimetry (DSC) and infrared thermal imaging camera. The DSC analysis showed that the exothermic heat released during the polymerization of PCLDMA increased with increasing concentrations of BPO/DMT. Formulation with 0.5 wt% BMO/DMT showed <4 cal/g exothermic heat release **(Figure 2B)**. To provide context, many common dietary carbohydrates and proteins provide about 4 cal/g of energy.^33^ This indicates that the ISCD cross-linking reaction is relatively mild and releases minimal heat. Additionally, monitoring the temperature changes during the polymerization process *via* an infrared thermal imaging camera showed a temperature of 20.9 °C, confirming the absence of localized heating **(Figure S3)**. These findings alleviate concerns about potential thermal tissue damage upon ISCD injection.

To assess the ability of ISCD for ultra-long-term release of hydrophilic drugs, we encapsulated TAF – a hydrophilic anti-retroviral with a water solubility of 5.63 mg/mL. We first assessed the injectability of the pre-polymer mixture with and without TAF (90 mg/mL) by determining their viscosities and calculating the injection force using the Hagen-Pouiselle equation^34^ under conditions specified in **Figure 2C** for a 23 gauge needle. The calculated injection forces for the pre-polymer mixture with and without TAF were 64.0 ± 3.7 N and 44.4 ± 7.8 N respectively, indicating an increase in viscosity and injection force with drug encapsulation **(Figure 2D,E)**. Importantly, both compositions exhibited injection force values below 80 N, which is considered the maximum acceptable injection force for most people,^34^ confirming the injectability of these formulations.

*In vitro*, ISCD demonstrated sustained release of TAF, with ∼50% cumulative release over 250 days. This was observed for ISCD depots comprising either PCLDMA or PCLTMA **(Figure 2F)**. TAF release kinetics was similar across a range of drug loadings (30-150 mg/mL) **(Figure 2G)**. This indicates that the rate of drug release is independent of the initial amount of drug loaded, but the total amount of drug released ultimately scales proportionally with the initial loading **(Figure S4)**.

To function effectively, LAI depots need to maintain their mechanical integrity at the injection site. Therefore, we used a mechanical tester to understand the mechanical properties of ISCD by performing compression testing. ISCD with and without TAF (90 mg/ml) were prepared in a cylindrical mold made of polydimethylsiloxane (PDMS). The compressive stress moduli for ISCDs with and without TAF were 68.1 ± 10.3 MPa and 67.2 ± 4.0 MPa, respectively, and the yield stresses were 24.2 ± 1.1 MPa and 25.9 ± 1.5 MPa, respectively **(Figure 2H,I)**. These mechanical properties closely align with those observed in clinically used solid implantable devices^13, 35^, confirming the structural rigidity of ISCD. Importantly, the inclusion of TAF does not compromise the structural integrity of the delivery system.

We also demonstrated that ISCD can be molded and crosslinked *ex vivo* into various shapes, including cylinders, pipes, and disks, using PDMS molds. These forms maintain a similar drug release profile to the injectable version **(Figure 2J, S5)**, providing a versatile platform for both injectable and implantable drug delivery.

### Tailoring the drug release kinetics and degradation of ISCD

Having demonstrated the ultra-long-term release of TAF from ISCD, we aimed to identify design parameters that can tailor the polymer network to fine-tune the drug release kinetics. TAF concentration in ISCD was maintained at 90 mg/ml for all these experiments. We hypothesized that modulating intrinsic factors of ISCD, including the molecular weight of PCLDMA, and the concentrations of BPO and DMT, could influence the cross-linking density of ISCD, thereby impacting drug release. To evaluate the impact of polymer molecular weight on drug release, TAF-loaded ISCDs (90 mg/mL TAF) were formulated using PCLDMA with two distinct molecular weights (630 Da and 2100 Da) **(Figure 3A)**. As expected, depots with higher molecular weight polymer showed faster release of TAF. Pevious reports have demonstrated that polymer molecular weight of LAIs impacts crosslinking density, which in turn affects drug release.^36, 37^ Similarly, release profiles of TAF from ISCDs with varying concentrations of BPO/DMT ranging between 0.1 to 0.4 wt% demonstrated that an increase in the concentrations of the initiator and accelerator reduces the burst release and the overall release rate **(Figure 3B)**. To confirm if the effect of BMO/DMT concentration on drug release kinetics is attributed to the changes in the cross-linking density of the depot, we quantified the degree of cross-linking of different ISCD depots using a gravimetric approach. The depots were incubated in benzyl alcohol for a week to determine swelling and the Flory-Rehner equation^38^ was used to estimate the degree of cross-linking based on the swelling data. As expected, reducing BPO/DMT concentrations resulted in a reduction in the cross-linking density **(Figure 3C)** and an increase in swelling percentage **(Figure S6)**, confirming our hypothesis.

**Figure 3.**
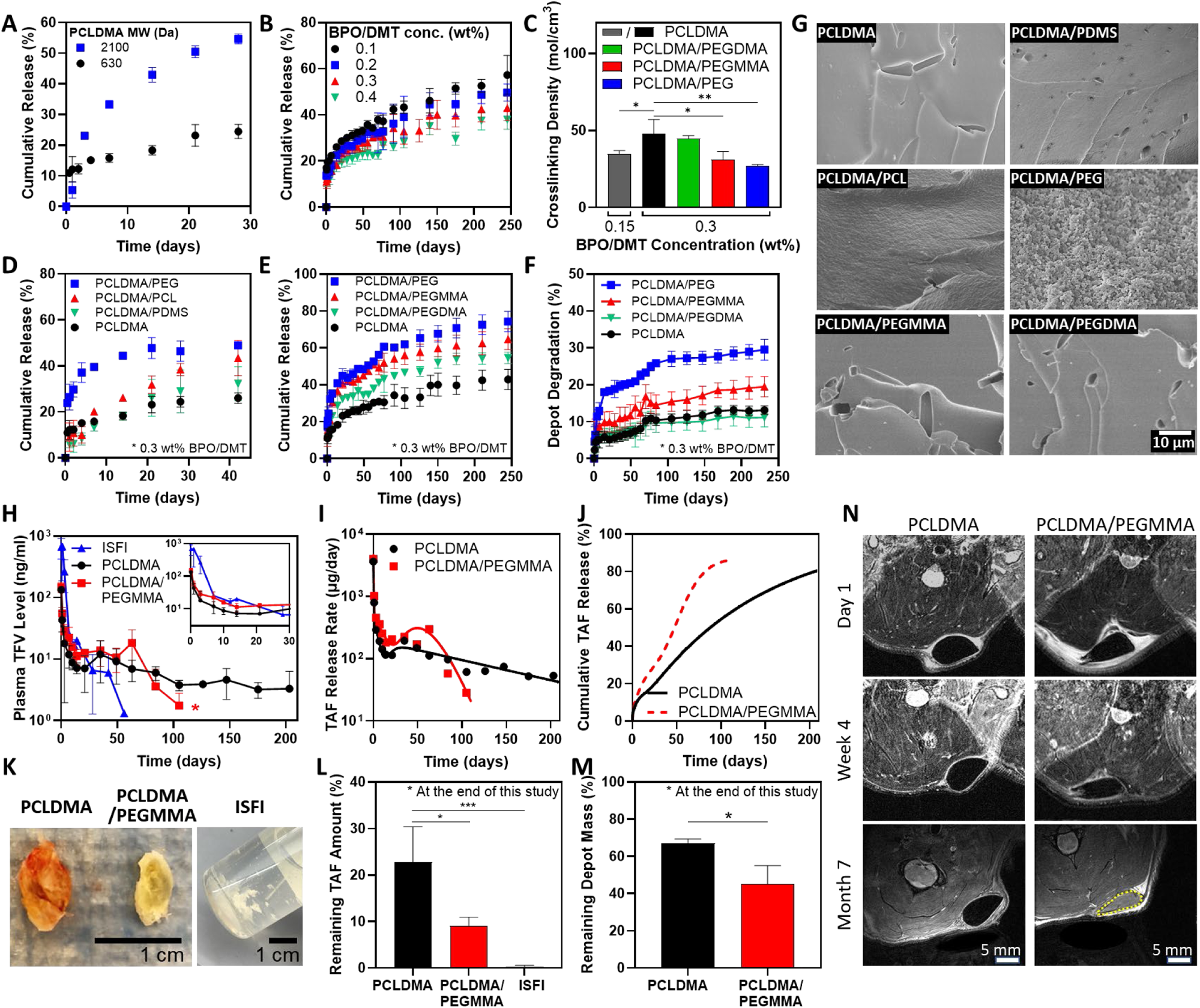
Tailoring the drug release kinetics and degradation of ISCDs *in vitro* and *in vivo*. **A.** *In vitro* release profiles of TAF in PBS (37°C) from ISCDs prepared with PCLDMA of different molecular weights (630 Da and 2100 Da). **B.** *In vitro* release profiles of TAF in PBS (37°C) from ISCDs prepared with varying BPO/DMT concentrations. **C.** Crosslinking density of unmodified ISCD prepared with different concentrations of BMP/DMT and ISCD containing different external polymer additives (25 wt%) (*P<0.05 and **P<0.01). **D.** *In vitro* release profiles of TAF in PBS (37°C) from unmodified ISCD (prepared using PCLDMA only) or ISCD containing 25 wt% of an external polymer additive (PEG, PCL, or PDMS) alongside PCLDMA. **E.** *In vitro* release profiles of TAF and **F.** and percentage depot degradation in PBS (37°C) for unmodified ISCD or ISCD containing 25 wt% of PEG with varying degrees of methacrylation. **G.** SEM images of unmodified ISCD or ISCD containing different external polymer additives (25 wt%) showing the cross-section of depot structure at week 1 post-incubation in PBS (37°C). **H.** Plasma level of TFV in rats injected with 500 µl of TAF-loaded ISFI (control) or TAF-loaded unmodified ISCDs of ISCD containing 25 wt% PEGMMA. All depots were loaded with 90 mg/mL of TAF. The inset shows plasma levels up to day 30. (*P<0.05 for the overall comparison of plasma levels of the two ISCDs over the entire study duration). **I.** *In vivo* daily release rate and **J.** cumulative release of TAF from unmodified ISCD or ISCD containing 25 wt% of PEGMMA, as determined by PK modeling. **K.** Camera images of TAF-loaded ISFI or TAF-loaded unmodified ISCD or ISCD containing 25 wt% of PEGMMA, retrieved from rats at month 7 post-injection, and **L.** Remaining TAF amount in the depots (*P<0.05 and ***P<0.001) and **M.** Remaining mass of the depot (*P<0.05). Due to the disintegration of ISFI within the animal, the remaining mass of the ISFI depots could not be measured. **N.** MRI images of subcutaneously injected unmodified ISCD or ISCD containing 25 wt% PEGMMA at different time points. Data in A, B, C, E, F, and G are presented as mean ± standard deviation (n=3, experiments performed at least twice). Data in H, L, and M are presented as mean ± standard deviation of technical repeats (n=3, experiment performed twice). Data in I and J present predictions from PK modeling of the average plasma levels of TFV obtained experimentally. *P*-value in H was determined using two-way ANOVA with Bonferroni correction, with time and different ISCD formulations as the two variables. The *P*-value in C and L was determined using one-way ANOVA with Tukey’s post hoc analysis. The *P*-value in M was determined using Student’s *t*-test.

Next, we hypothesized that incorporating hydrophilic polymer additives along with PCLDMA would modulate the hydrophilicity of the depot, influencing drug release kinetics. To test this, we added 25 wt% of different polymer additives with varying hydrophilicity but similar molecular weights: polyethylene glycol (PEG, MW 500), PCL-diol (MW 530), and poly(dimethylsiloxane) (PDMS, MW 500). PEG’s oxygen-rich polymer chain promotes strong interactions with water, making it highly hydrophilic,^39^ while PCL-diol, with hydrocarbon chains and polar ester groups, displays moderate hydrophobicity.^40^ PDMS, with its silicone-based structure and methyl-covered surface, is the most hydrophobic among the three polymers.^41^ We chose non-methacryalted polymers to ensure that the effects observed on the release kinetics are purely due to the variation in hydrophilicity. Relase profiles of TAF from these ISCDs were compared to the release from unmodified ISCD formulated with pristine PCLDMA. The data support a positive correlation between the hydrophilicity of the additive and the drug release rate from ISCDs. ISCDs containing PEG exhibited the highest cumulative release (46.5 ± 4.5%) at 42 days post-incubation **(Figure 3D, S7)**. This was nearly double the release observed for the unmodified ISCD (24.5 ± 2.3% on day 42). PCL-diol, with intermediate hydrophobicity, also led to a significantly increased release (38.4 ± 2.7%) compared to the unmodified ISCD. Conversely, PDMS, the most hydrophobic additive, showed minimal impact on drug release. These findings suggest that incorporating hydrophilic additives into the polymer network of ISCDs can enhance the drug release rate.

Building upon our investigation into the influence of external polymer additives, we sought to explore if methacrylation of the hydrophilic polymer additives and the degree of methacrylation can further impact drug release. To study this, we incorporated 25 wt% of non methacrylated PEG (PEG), singly-methacrylated PEG (polyethylene glycol monomethyl ether mono-methacrylate, PEGMMA), or double-methacrylated PEG (polyethylene glycol dimethacrylate, PEGDMA) along with PCLDMA. All PEG derivatives had the same molecular weight of 500. Increasing the degree of methacrylation reduced both burst release and the overall release rate **(Figure 3E)**. By 240 days, ISCDs with PEG, PEGMMA, and PEGDMA showed cumulative releases of 72.6 ± 5.3 %, 62.5 ± 6.3 %, and 54.2 ± 3.1 %, respectively, which were all significantly higher than the cumulative release of 42.4 ± 5.5 % observed for the unmodified ISCD **(Figure S8A)**.

To elucidate the mechanism of increased drug release due to the incorporation of polymer additives, we performed scanning electron microscopy (SEM) based qualitative assessment of depots at week 1 post-incubation in PBS at 37°C and also determined their cross-linking density. Depot morphology aligned well with the observed drug release behavior **(Figure 3G)**. ISCD containing PEG, the most hydrophilic additive, displayed a highly porous surface compared to the control depot consisting of PCLDMA alone, which showed a densely packed smooth surface. This suggests erosion of PEG-containing ISCD, which could be attributed to the efflux of the uncrosslinked PEG polymer due to its hydrophilicity, leading to enlarged pores, and hence faster drug release than the unmodified ISCD. ISCD with PCL-diol, exhibiting an intermediate hydrophilic character, also showed mild erosion with surface roughness, which explains faster drug release than the unmodified ISCD. Conversely, ISCD with PDMS, the most hydrophobic additive, displayed minimal erosion and a smooth surface, consistent with minimal difference in drug release compared to the unmodified ISCD. Interestingly, PEG-containing ISCD depots also showed significantly lower cross-linking density and swelling percentage compared to the unmodified ISCD **(Figure 3C, S6)**. These observations support the conclusion that incorporating non-crosslinkable hydrophilic polymers as additives can significantly influence the drug release profile from ISCD by decreasing the cross-linking density and promoting erosion of the depot. Notably, unlike non-methacrylated PEG-containing ISCD, SEM images of PEGMMA or PEGDMA-containing ISCD didn’t show any signs of depot erosion **(Figure 3G)**, despite their higher TAF release compared to the unmodified ISCD **(Figure 3E)**. However, PEGMMA-containing ISCD showed a significantly lower cross-linking density and higher swelling percentage compared to the unmodified ISCD **(Figure 3C, S6)**. PEGDMA-containing ISCD on the other hand did not show significant changes in cross-linking density or swelling compared to the unmodified ISCD. These findings are consistent with the drug release kinetics, which was significantly faster for PEGMMA-containing depots compared to the ones with PEGDMA, and suggest that the increase in drug release observed with methacrylated hydrophilic polymers is correlated to their ability to reduce the cross-linking density of the network. Significantly increased drug release for PEGDMA-containing ISCD compared to the unmodified ISCD could be attributed to the inherent hydrophilic nature of PEGDMA, which can increase the overall hydrophilicity of the network, thereby increasing water permeability, leading to faster drug diffusion. The absence of depot erosion for PEGMMA and PEGDMA-containing ISCDs can be largely attributed to the ability of PEGMMA and PEGDMA to cross-link within the polymer network. Overall, our data shows that the incorporation of hydrophilic additives in ISCD can enhance drug release by promoting depot erosion, modulating the cross-linking density, or by simply increasing the overall hydrophilicity of the network.

The *in vitro* degradation of ISCD was also found to be dependent on the cross-linking density. Degradation was assessed by monitoring mass changes of ISCDs over time while incubated in PBS at 37 °C. After seven months, ISCDs with PEG and PEGMMA exhibited significantly faster degradation, reaching 28.9 ± 2.0% and 19.1 ± 3.0%, respectively, compared to the unmodified ISCD, 12.9 ± 1.3 % **(Figure 3F, S8B)**. In contrast, ISCD with PEGDMA showed a similar degradation profile as the control ISCD, reaching 11.1 ± 2.6 % at 7 months post-incubation, which is consistent with their similar cross-linking densities.

### Pharmacokinetics and *in vivo* degradation of ISCD

After demonstrating the ultra-long-term release of TAF from ISCD *in vitro*, we aimed to evaluate the pharmacokinetics (PK) of the TAF-loaded ISCD system *in vivo*. Our *in vitro* TAF release data showed faster drug release from PEGMMA-containing ISCD compared to unmodified ISCD. Based on these findings, we selected two ISCD compositions for the *in vivo* study: unmodified ISCD with pristine PCLDMA and ISCD with 25 wt% PEGMMA. Our goal was to confirm if the release tunability observed *in vitro* would also be evident *in vivo*.

We subcutaneously injected 500 µL of TAF-loaded ISCDs (90 mg/mL TAF) in rats and used a PLGA-based ISFI formulation as control. It should be noted that TAF is highly unstable in rodent plasma, and rapidly converts to tenofovir (TFV) due to high levels of plasma esterases expressed in rodent species which lead to hydrolytic cleavage of TAF.^14, 43^ Because of this reason, we couldn’t detect TAF levels in rat plasma, and therefore measured TFV levels to assess the PK. The *in vivo* release rate (µg/day) from ISCD formulations was determined using the area function method^42^ by comparing the ISCD PK data (**Figure 3H**) with intravenous (IV) PK data of a single bolus of TAF in rats. Additionally, we performed compartmental PK modeling to quantitatively characterize the *in vivo* PK profiles of ISCDs **(Figures S9-10, Tables 1, S2)**. This involved integrating an appropriate absorption kinetic model describing the *in vivo* release rate of ISCD (**Figure 3I**) with a two-compartment disposition kinetics model describing the IV PK data (**Figure S10A**).

Burst release, as characterized by the peak plasma TFV concentration (C_max_) at 4 hours post-injection was almost 5-fold lower for both unmodified ISCD (133.7 ± 12.7 ng/mL) and PEGMMA-containing ISCD (146.7 ± 27.3 ng/mL), as compared to the ISFI formulation (668.0 ± 246.0 ng/mL) **(Figure 3H)**. Following a minimal burst release (<6% of the total drug dose; **Table 1**), plasma TFV levels of rats injected with unmodified ISCDs reached less than 10 ng/mL within 10 days and maintained a sustained level of 1-10 ng/mL for at least 210 days (7 months). Based on our PK analysis, the unmodified ISCD maintained a steady daily release rate of 50-100 µg/day for at least 7 months **(Figure 3I)**, translating to a cumulative drug release of 81% at 7 months **(Figure 3J)**. PEGMMA-containing ISCD displayed a distinct release profile compared to unmodified ISCD, characterized by a higher release rate and shorter duration **(Figure 3H-J)**. After a minimal initial burst (<7% of the total dose), plasma TFV levels peaked below 20 ng/mL by day 10 and remained between 10-20 ng/mL until day 63. This was followed by a rapid decline, reaching undetectable levels (<1 ng/mL) after 120 days (4 months). Notably, PCLDMA/PEGMMA released TFV at a significantly higher rate (∼200 µg/day) for 2 months compared to PCLDMA **(Figure 3I)**, resulting in a cumulative release of 86% at 3 months **(Figure 3J)**. These PK data are consistent with the trends observed in our *in vitro* release kinetics study. Importantly, both ISCD formulations demonstrated significantly longer sustained release compared to the conventional ISFI system, which exhibited a high initial burst release followed by a rapid decline in plasma TFV levels, falling below the detection limit after 2 months.

**Table 1.** Drug release kinetic parameters for drug-loaded ISCD formulations.

| Drug Release phase | TAF-loaded PCLDMA ISCD |  |  | TAF-loaded PCLDMA/PEGMMA ISCD |  |  |
| --- | --- | --- | --- | --- | --- | --- |
|  | Rate constant (1/day) | Cumulative release (%) | Time delay (day) | Rate constant (1/day) | Cumulative release (%) | Time delay (day) |
| Initial | 2.47 | 5.8 |  | 2.52 | 6.8 |  |
| Intermediate | 0.178 | 9.2 | 1.5 | 0.0643 | 27.1 | 1.0 |
| Sustained | 0.00744 | 65<sup>#</sup> | 18.1 | 0.128 | 52.1<sup>#</sup> | 50.8 |
| Drug Release phase | TAC-loaded PCLDMA ISCD |  |  | NAL-loaded PCLDMA ISCD |  |  |
|  | Rate constant (1/day) | Cumulative release (%) | Time delay (day) | Rate constant (1/day) | Cumulative release (%) | Time delay (day) |
| Initial | 0.962 | 8.7 |  | 0.372 | 27.4 |  |
| Intermediate | 0.122 | 15.6 | 3.7 | 0.0346 | 15.4 | 11.8 |
| Sustained | 0.0134 | 10.7<sup>#</sup> | 68.3 | 0.0181 | 54.2<sup>#</sup> | 70.1 |

<sup>#</sup>Calculated as %Total drug mass released in vivo by the end of the study -%Release by initial and intermediate phases

At the end of the study, we retrieved the depots from euthanized rats **(Figure 3K)**, and the remaining drug load was quantified using CHN analysis **(Figure 3L)**. Significantly higher amounts of remaining TAF were observed in the unmodified ISCD compared to PEGMMA-containing ISCD (22.8 ± 3.4% for PCLDMA and 9.1 ± 1.8% for PCLDMA/PEGMMA). Notably, explants from ISFIs at month 2 post-injection exhibited minimal drug remaining (0.26 ± 0.20%) **(Figure S11A)**. ISCDs could be extracted from tissue as a single, easily removable solid depot **(Figure 3K)**, demonstrating superior retrievability, whereas ISFIs were observed as fragile solids, prone to fracture and difficult to extract from the tissue **(Figure S11B)**. We also measured the mass of the remaining depots to assess the biodegradation of ISCD. The remaining mass of the extracted depots was found to be 67.1 ± 2.8% for the unmodified ISCD and 45.2 ± 12.1% for PEGMMA-containing ISCD, which correspond to 33% and 55% *in vivo* degradation, respectively **(Figure 3M)**. The trend is consistent with the *in vitro* degradation data and confirms that the addition of PEGMMA increases the degradation of ISCD. We also used magnetic resonance imaging (MRI) to evaluate the long-term morphological changes of ISCDs *in vivo* and to study the interactions between depots and host tissues at day 1, month 1, and month 7 after injection **(Figure 3N)**. We confirmed that both unmodified and PEGMMA-containing ISCDs did not migrate to other sites or cause adverse tissue reactions. The size of the depots decreased over time, indicating biodegradation, but both depots maintained good structural integrity. Interestingly, at month 7 post-injection, we found that the PEGMMA-containing ISCD appeared white on MRI images, as did the tissue surrounding the depot, indicating that they had a higher water content than before, while the PCLDMA-only depots remained dark. Since PEGMMA is hydrophilic, higher water content could be attributed to the penetration of water into the depots, as the polymer network loosened over time.

It was critical to confirm if TAF encapsulated within the ISCDs didn’t degrade into TFV and maintained its chemical structure prior to release. To address this, we employed high-performance liquid chromatography (HPLC) analysis of explanted depots retrieved at 7 months post-injection. The HPLC data of the dissolved explants exhibited a single peak eluting at the same retention time as the freshly prepared TAF solution **(Figure S12)**. This confirms the absence of degradation products within the retrieved depots, indicating that TAF remains stable during the crosslinking process and within the depot under *in vivo* conditions prior to its release.

### Biocompatibility and safety of ISCD

*In vivo* biocompatibility is essential for LAIs to minimize adverse reactions and inflammation at the injection site, ensuring patient safety and efficacy.^6, 44^ We evaluated the biocompatibility of ISCD consisting of pristine PCLDMA without any drug. Depots were injected on day 0, followed by histological and immunohistochemistry (IHC) analysis of the local tissue explanted at week 1, week 4, and month 7 post-injection. The H&E staining and IHC analysis revealed an initial inflammatory response at week 1, characterized by the presence of immune cells, including CD3-positive T cells and CD68-positive macrophages, around the depot **(Figure 4A-C)**. However, by week 4, there was a substantial decrease in the number of immune cells around the depot **(Figure 4B)**, accompanied by a significant reduction in CD3 and CD68 positive populations **(Figure 4C)**, a trend that persisted at month 7, resulting in negligible immune cells present around the depot. Importantly, the tissue samples exhibited no signs of fibrosis, a major complication associated with implant failure and typically identified by excessive collagen deposition.^45^ These findings indicate successful integration of the depot with the surrounding tissue.

**Figure 4.**
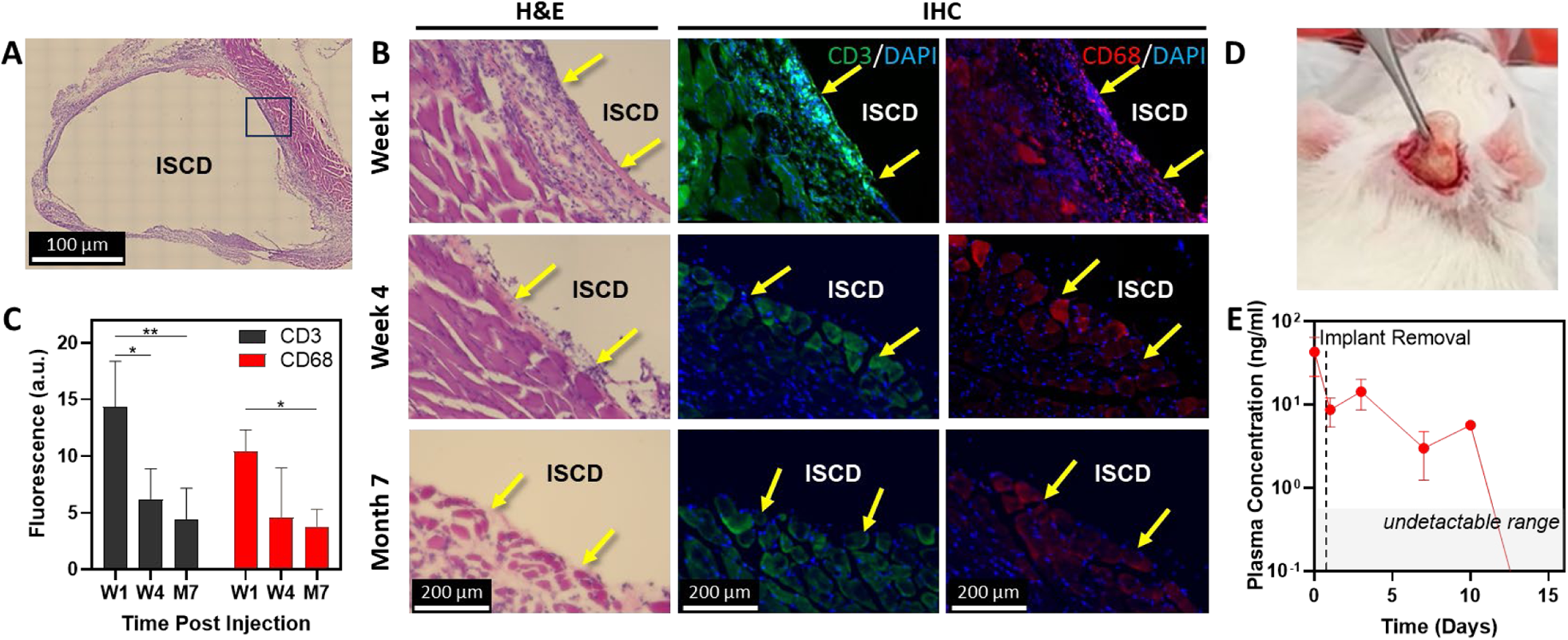
*In vivo* biocompatibility and safety of ISCD. **A.** Representative image (10X magnification) of an H&E stained section of local tissue, explanted with the ISCD depot one week after subcutaneous injection of 500 µl PCLDMA-based ISCD in rats. **B.** Left side shows high magnification (20X) representative images of H&E-stained sections of local tissue, explanted with the ISCD depot at different time points. Yellow arrows show inflammatory cells. The right side shows representative immunofluorescence images of local tissue sections explanted at different time points and stained against CD3 (green) and CD68 (red) markers to visualize T cells and macrophages, respectively. Yellow arrows show cells positive for CD3 or CD68. **C.** Fluorescence intensity quantified for CD3 and CD68 immunofluorescence. (*P<0.05, **P<0.01). **D.** Camera image taken during the procedure of retrieving ISCD from a rat, showing safe retrievability *via* a small incision. **E.** Plasma levels of TFV following ISCD removal. Data in C and E are presented as mean ± standard deviation of technical repeats (n=3). The *P*-value in C was determined using one-way ANOVA with Tukey’s post hoc analysis.

Although ISCD is biocompatible, the drug delivered *via* ISCD may exhibit adverse effects, necessitating prompt depot retrieval. In such cases, a rapid decrease in plasma drug levels following depot removal would be crucial to mitigate side effects. To confirm this, we subcutaneously administered TAF-loaded ISCD to rats and retrieved the depots two weeks later through a small incision near the injection site on the skin **(Figure 4D)**. Plasma TFV levels were monitored after retrieval. Following the depot removal, plasma TFV concentrations showed an exponential decline, decreasing more than four-fold within a day to less than 10 ng/mL **(Figure 4E)**. By day 10 post-retrieval, TFV plasma concentrations had dropped below the limit of detection (1 ng/ml) for two out of three rats, and the plasma level went below detection for all three rats by day 14, confirming the safety of ISCD.

### Versatility of ISCD for a wide range of therapeutics and combination therapies

Next, we wanted to understand the versatility of the ISCD platform for delivering a broad spectrum of hydrophilic drugs with water solubilities in the range of 3-112 mg/mL. We chose drugs representing diverse therapeutic classes: antiretrovirals, including emtricitabine (FTC), abacavir (ABC) and lamivudine (LAM), an opiate antagonist, naltrexone (NAL), and antibiotics, including vancomycin (VAN) and amoxicillin (AMX). We also evaluated a hydrophobic drug – tacrolimus – a clinically used immunosuppressant. All drugs were encapsulated at a concentration of 90 mg/mL for direct comparison of the release profiles between different drugs. *In vitro*, ISCD demonstrated ultra-long-term release of all the therapeutics over at least 150-360 days **(Figure 5A).** Our analysis revealed a positive correlation between a drug’s hydrophilicity and its cumulative release on day 1. Drugs with higher water solubility (more hydrophilic) showed a higher initial release compared to drugs with lower water solubility **(Figure 5B, Table S1)**. This can be explained by the stronger interaction between hydrophobic drugs and the hydrophobic polymer backbone of PCLDMA. However, ISCD successfully minimized the overall burst release for all the drugs. Even for the drugs with water solubilities as high as 100 mg/mL, such as NAL and FTC, day 1 cumulative release was ∼20%, which is significantly lower than the cumulative release reported for hydrophilic drugs from injectable systems developed previously.^46, 47^

**Figure 5.**
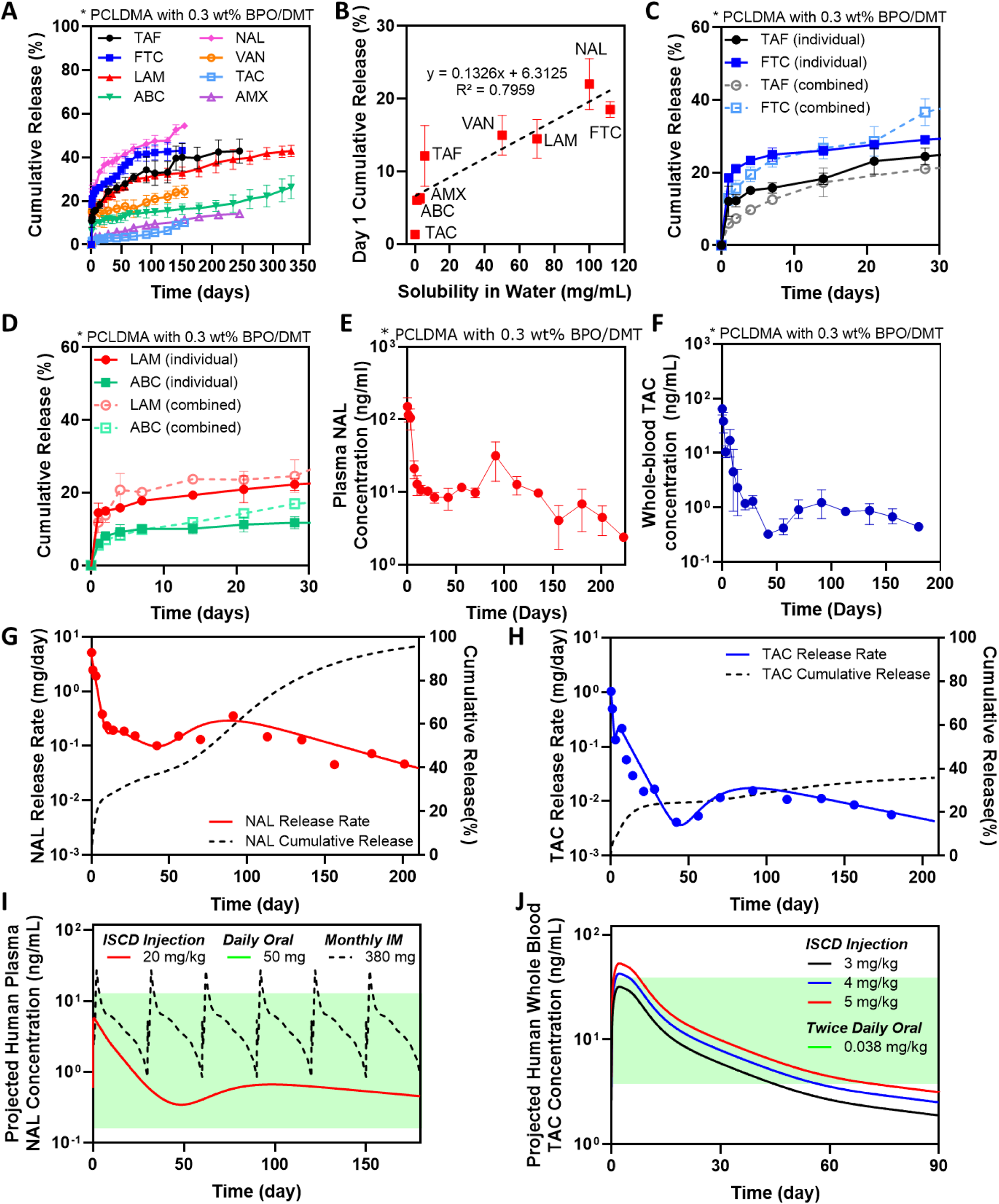
The versatility of the ISCD platform and human PK prediction. **A.** *In vitro* release profile of different drugs with varying water solubilities encapsulated into the ISCD platform. The release was studied in PBS (37°C). **B.** Correlation of cumulative release at day 1 with different drugs with varying water. **C.** *In vitro* release profile of TAF and FTC, when loaded into the ISCD platform individually versus in combination. The release was studied in PBS (37°C). **D.** *In vitro* release profile of ABC and LAM, when loaded into the ISCD platform individually versus in combination. The release was studied in PBS (37°C). **E.** Plasma concentration of NAL in rats subcutaneously injected with 500 µl of NAL-loaded ISCD (45 mg/ml NAL). **F.** Whole blood concentration of TAC in rats subcutaneously injected with 500 µl of TAC-loaded ISCD (28 mg/ml TAC). *In vivo* daily release rate and cumulative release profile of **G.** NAL and **H.** TAC, as predicted by PK modeling of systemic drug levels in rats, following subcutaneous injection of 500 µl of NAL- or TAC-loaded ISCD. **I.** Convolution analysis-based prediction of human PK of a single subcutaneous dose of NAL-loaded ISCD in comparison to clinically established PK profile of once-daily oral dose of NAL (green region), and once monthly intra-muscular injection – Vivitrol® (purple lines). **J.** Convolution analysis-based prediction of human PK of a single subcutaneous dose of TAC-loaded ISCD (at different dosages) in comparison to clinically established PK profile of twice-daily oral doses of TAC (green region). Data in A-D are presented as mean ± standard deviation (n=3, experiments performed at least twice). Data in E and F are presented as mean ± standard deviation of technical repeats (n=3). Data in G and H present predictions from PK modeling of the average plasma level of TAC and NAL obtained experimentally. Data in I and J present human PK prediction based on convolution analysis of experimentally obtained PK data of TAC and NAL in rats.

We also demonstrated the potential of ISCD for co-delivery of combination therapies. We encapsulated two clinically used combination regimens for HIV therapy: Epzicom (FTC and LAM) and Descovy (ABC and TAF). We compared the release kinetics of these drugs from ISCD when encapsulated individually versus in combination. Release profiles of drugs encapsulated individually were almost identical to those drugs encapsulated in combination **(Figure 5C,D)**. This suggests that ISCD can be formulated to co-deliver multiple drugs while maintaining their independent release characteristics, making it a promising platform for long-acting combination therapy.

Building on the *in vitro* demonstration of ISCD’s versatility, we conducted a PK study to validate the ultra-long-term release of drugs with different water solubilities *in vivo*. We selected two drugs with contrasting water solubilities compared to TAF, which showed ultra-long-term release *in vivo*. These included TAC, which has lower water solubility than TAF, and NAL, a drug with higher water solubility than TAF **(Table S1)**. Rats were subcutaneously injected with 500 µL of PCLDMA ISCDs containing either 45 mg/mL of TAC or 90 mg/mL of NAL. Since our *in vitro* data showed a faster release of NAL compared to TAC, we used a higher concentration of NAL than TAC. Blood samples were collected at intervals up to 6 or 7 months post-injection to analyze whole-blood concentration of TAC and plasma concentration of NAL. Similar to TAF, we performed PK analysis for TAC and NAL-loaded ISCDs by utilizing both non-compartmental and compartmental methods. For compartment modeling of the PK profiles **(Figures S9,10, Table S2)**, we established the disposition kinetics by two-compartmental analysis of PK data obtained from experimental bolus intravenous injection of TAC and NAL in rats. Consistent with the *in vitro* release data, ISCD showed a higher initial burst release for NAL *in vivo* (27%) compared to TAC (5.8%) **(Figure 5G,H**, **Table 1)**, with C_max_ values of 64.9 ± 13.0 ng/mL and 150.7 ± 80.5 ng/mL for NAL and TAC, respectively. Following the initial burst, the ISCD established sustained drug release. As expected based on the water solubilities of NAL and TAC, the systemic concentration of NAL was maintained within a range of 5-15 ng/mL, while TAC concentrations were lower, ranging from 0.5-1.5 ng/mL. This translated to a cumulative release of approximately 92% for NAL and 35% for TAC in 6 months **(Figure 5G,H)**. These findings demonstrate the versatility of the ISCD platform to enable ultra-long-term release of drugs with a wide range of water solubility. The data also establish a clear relationship between a drug’s hydrophilicity and its PK, when delivered using an ISCD system.

Convolution analysis was utilized to predict the human PK of NAL- and TAC-loaded ISCDs. Human disposition kinetics information was obtained by the analysis of previously reported human PK data for NAL IV bolus^48^ and TAC administered orally^49^ **(Table S3)**. These human disposition kinetics were then integrated with the release function derived from the PK analysis of NAL and TAC-loaded ISCDs in rats, assuming simple allometry on release rate constants between species. The predicted human PK profiles for NAL and TAC from ISCDs (**Figure 5I-J**) were compared with the PK of their clinically available formulations, including oral TAC^49^, oral NAL^48^, and an intramuscular (IM) LAI of NAL (Vivitrol®)^50^. Notably, a single dose of NAL-loaded ISCD, when injected at 20 mg/kg is predicted to maintain prolonged steady plasma concentrations for at least 6 months, well within the established peak and trough levels observed with once-daily oral doses of NAL tablets at 50 mg **(Figure 5I)**.^51^ Compared to Vivitrol®, the projected PK profile of NAL-loaded ISCD showed 4.6-fold lower burst release.^50^ Interestingly, while Vivitrol®, with a NAL dose of 380 mg, requires a once-monthly injection, the ISCD, with only ∼4 times the dose of NAL compared to Vivitrol®, maintains steady plasma concentrations for at least 6 months, suggesting the potential to improve the dosing schedule from once a month to once every 6 months. Similarly, the predicted PK profile of TAC from 3, 4 and 5 mg/kg TAC-loaded ISCDs showed sustained release for at least 6 months **(Figure S13)**, with blood levels maintained within the therapeutic concentration range of twice-daily oral doses of TAC capsules (0.038 mg/kg) for 2-3 months **(Figure 5J)**.^49^ Overall, our data clearly suggest that ISCD has the potential to enable ultra-long-term release of both hydrophilic and hydrophobic drugs in humans.

## Discussion

We present ISCD, an *in situ* cross-linking, biocompatible, and tunable LAI platform designed for ultra-long-term delivery of hydrophilic therapeutics. ISCD is a versatile platform that demonstrated sustained release of multiple hydrophilic drugs with varying water solubilities as high as 100 mg/mL, and from diverse classes of therapeutics, including antiretrovirals, opioid antagonists, and antibiotics. A single subcutaneous injection of TAF-loaded ISCD in rats resulted in significantly prolonged drug release (>7 months) with lower burst release compared to the conventional ISFI system. Prolonged release in rats was also observed for NAL-loaded ISCD, which translated to a predicted 6-month release duration in humans based on PK modeling. This represents a significant improvement over Vivitrol®, a clinically used LAI for NAL, which requires monthly injections and has a higher initial burst. ISCD also demonstrates ultra-long-term release of TAC-a hydrophobic drug.

Hydrophobic drugs have been successfully formulated into LAIs that achieve ultra-long-term release over several months. Examples include the bimonthly LAIs for HIV drugs cabotegravir and rilpivirine (APRETUDE® and CABENUVA®), and the recently approved lenacapavir as once every 6 months (SUNLENCA®). Leuprolide acetate, a synthetic hormone used for various conditions including prostate cancer,is available as a once-every-six-months LAI (Eligard®). In contrast, achieving ultra-long-term delivery of hydrophilic drugs, which constitue a substantial portion of therapeutics, remains a significant challenge with LAIs. Currently available LAIs for hydrophilic drugs offer only limited release duration, up to a month, and often result in high burst release.^22^ Examples include Zoladex®, a once-monthly injection of goserelin acetate for prostate cancer; Nutropin® Depot, a once- or twice-monthly injection of somatropin for growth failure; Sublocade®, a monthly buprenorphine injection for substance use disorder (SUD); and Vivitrol®, a monthly naltrexone injection for SUD. Additionally, current LAIs are limited to delivering single drugs and have not demonstrated the capability of co-delivery of multiple drugs, in which is crucial for combination therapies, which are the gold standard for treating conditions like HIV. These limitations in LAI technology present a significant barrier to leveraging the full therapeutic potential of hydrophilic drugs, and research dedicated to developing improved LAI platforms for these drugs has been lacking. To our knowledge, this is the first report demonstrating the ultra-long-term release of hydrophilic drugs, either individually or in combination, using an LAI.

Our approach offers several advantages over existing methods for long-term delivery of hydrophilic drugs. ISCD offers simple and minimally invasive administration. While implantable devices have demonstrated ultra-long-term delivery of hydrophilic drugs, they require invasive, time-consuming medical procedures for insertion, which can be challenging, especially in low-resource settings and in low- to middle-income countries.^26, 27^ For instance, a nanofluidic implant designed for the ultra-long-term delivery of anti-retroviral drugs, including TAF, necessitates a surgical procedure for implant insertion.^52, 53^ This involves making a small incision to create a subcutaneous pocket for the implant, followed by sealing the wound with sutures or surgical adhesive under local anesthesia. In contrast, ISCD requires simple subcutaneous or intramuscular injection of the pre-polymer-drug mixture. The final clinical product could be a simple two-component system consisting of 1) a pre-polymer/drug mixture and 2) BPO/DMT mixture. This could be either injected using a double barrel syringe or mixed in a vial a few minutes before the injection and injected usual a regular syringe. Another concern for certain implants, such as the nanofluidic implant described above, is that they are non-biodegradable, and require surgical removal after the drug is fully released. ISCD is degradable, and doesn’t require surgical removal. Furthermore, ISCD circumvents the limitations of solvent-based LAI formulations (ISFI) such as Sublocade®, which experience significant initial drug loss due to solvent efflux,^54^ thereby minimizing the overall release duration. ISCD is a solvent-free system, attributed to the ultra-low molecular weight of the pre-polymer PCLDMA, which helps minimize the burst release. In our study, we demonstrated that TAF-loaded ISCD exhibited a 10-fold lower initial burst release compared to TAF-loaded ISFI, resulting in a significantly prolonged drug release duration of 7 months with ISCD, compared to the 2-month release observed with ISFI. ISCD also exhibits significantly extended release profile compared to microsphere-based LAI approaches such as Vivitrol®. The porous matrix of microspheres promote significant water influx and efflux, resulting in a rapid drug release and a shorter duration. Our SEM data confirmed that ISCD forms a solid depot with dense polymer network, attributed to the ultra-low molecular weight PCLDMA, which limits the water influx and efflux, resulting in prolonged drug release. Previous studies in rats demonstrated that Vivitrol® at 50 mg/kg provided a one-month release of NAL^28^. In contrast, ISCD significantly extends the release duration to at least six months with only a three times higher dose than that of Vivitrol®. The predicted human PK also showed that ISCD can sustain steady plasma concentrations for at least 6 months, compared to only 1 month, as previously documented for Vivitrol®.^55^. Finally, ISCD exhibits excellent retrievability, as it can be easily explanted as a single solid depot, whereas ISFIs are prone to fragmentation during retrieval, as observed in our study.

Our study has several strengths. First, to achieve ultra-long term delivery of hydrophilic drugs, we employed a strategically designed yet simple approach. We utilized an ultra-low molecular weight PCL, a polymer that has been used in several FDA-approved products.^40^ The ultra-low molecular weight of PCL enabled ISCD to be solvent-free, while forming a dense hydrophobic matrix that minimizes water influx and efflux. To cross-link the methacrylated pre-polymer, we used BPO/DMT-based radical polymerization, a chemical reaction that has been employed in clinical bone cements for over 50 years.^56^ Thus, all components of ISCD, including PCL, BPO and DMT have a well-established history of clinical use. Second, we conducted in-depth studies to identify design parameters that can tailor the polymer network to tune the drug release kinetics and degradation of ISCD. To gain mechanistic understanding of how different external additives control the release kinetics, we performed rigorous analysis of cross-linking density using the Flory-Rehner equation and conducted SEM imaging to understand the contribution of surface morphology to the release kinetics. Third, we performed a comprehensive investigation exploring a wide range of hydrophilic drugs to demonstrate the platform’s versatility and understand the impact of drug hydrophilicity on release profiles. We studied the release of seven different hydrophilic drugs and one hydrophobic drug for up to 12 months *in vitro,* and validated the release of two hydrophilic and one hydrophobic drug for up to 7 months *in vivo*. We also demonstrated the potential of ISCD for co-delivery of multiple drugs for combination therapy. Fourth, we validated the PK model-based prediction of cumulative release of TAF from the ISCD by quantifying the total drug remaining in the depot using CHN analysis at the end of the study. The cumulative release predictions suggested 19% and 14% of TAF remaining for unmodified ISCD and PEGMMA-containing ISCD, respectively, which aligned well with the 22% and 9% remaining TAF quantified using CHN analysis. Minor discrepancies likely stem from the inherent limitations of the PK model and variability. Nonetheless, this close correspondence between experimental data and model predictions underscores the accuracy and utility of our PK model in describing drug release from ISCD. Finally, to establish translational feasibility of this platform, we performed human PK predictions for ISCD loaded with NAL and TAC, and demonstrated superior PK profiles and simplified dosing schedules compared to their clinically used formulations.

While this study establishes a foundation for the ISCD platform, there are limitations and several additional questions that future research needs to address. First, exploring diverse physicochemical properties of therapeutics beyond hydrophilicity will provide a more comprehensive understanding of ISCD release kinetics. While we found a strong correlation between the hydrophilicity of drugs and their release from ISCD, other factors, such as molecular weight and charge, also likely influence a drug’s release profile. Therefore, delving deeper into these interactions will be instrumental in optimizing the ISCD platform to accommodate a wider range of therapeutics. Second, while our *in vitro* data successfully demonstrated extended delivery of drug combinations with a single ISCD injection, future studies should validate this *in vivo*. Although we observed minimal interference between multiple drugs in ISCD, investigating the potential impact of complex biological processes on drug interactions *in vivo* will further elucidate ISCD’s performance in a living organism for combination therapy. Third, while human PK predictions for TAC-loaded ISCD indicated sustained drug release for at least six months, blood levels fell below the therapeutic window after 2-3 months. This could be attributed to the strong hydrophobicity of TAC combined with limited water influx/efflux in the ISCD, which can result in an ultra-slow release rate, as also observed from the PK analysis of the rat study. Future studies focusing on developing ISCD depots can leverage the design parameters to fine-tune the release kinetics and degradation of TAC-loaded ISCDs, ensuring blood levels are maintained above the therapeutic window for extended durations. Finally, to fully validate the clinical potential of the ISCD platform, it is essential to conduct PK and efficacy studies using clinically relevant large animal models, such as non-human primates. This will further validate the ISCD platform for rapid clinical adoption.

Taken together, our approach represents a paradigm shift in long-acting drug delivery, offering the remarkable capability of sustained release of both hydrophilic and hydrophobic drugs over unprecedented durations. This study also provides critical insights into optimizing drug release kinetics in ISCD and highlights the platform’s biocompatibility, retrievability, and ability to co-deliver multiple drugs. The platform promises to revolutionize healthcare by addressing the challenges of chronic diseases where consistent medication adherence is critical for therapeutic efficacy.

## Methods

### Synthesis of methacrylated polycaprolactone (PCL)

20 ml of 530 Da PCL-diol (Sigma-Aldrich) was mixed with 200 ml dichloromethane in a sealed nitrogen-filled flask chilled to 0°C. 33.8 mL of TEA (Sigma-Aldrich) was added to the flask using a syringe under stirring at 400 rpm, and 36 mL MAA (Sigma-Aldrich) was subsequently added dropwise using a syringe to have three molar eqivalent of TEA and MAA per mol of hydroxy group from PCL-diol.^57^ The reaction proceeded for 15 hours under stirring at 0°C. The reaction solvent was then removed under reduced pressure and reconstituted in 200 mL of ethyl acetate. The product was washed three times with each solvent: saturated aqueous sodium bicarbonate, 0.1 N hydrochloric acid (HCl), and aqueous brine in sequence. Following the washing steps, the product was dried with anhydrous sodium sulfate, filtered under vacuum, and solvent was removed under reduced pressure. The crude product was run through a pre-manufactured silica column (Teledyne Isco) using five column volumes of a mixture of 90% heptane/ 10% ethyl acetate, followed by five column volumes of pure ethyl acetate. Solvent was removed from the latter five fractions under reduced pressure to yield the purified product, PCLDMA, as a viscous yellow liquid. Substitution and purity were confirmed by ^1^H NMR.

### Preparation of pre-polymer mixture and ISCD depots for *in vitro* and *in vivo* samples

Pre-polymer solution, composed of methacrylated PCL (PCLDMA or PCLTMA) with or without external polymer additives (PEG, PEGMMA, PEGDMA, PCL, or PDMS), was mixed with 0.1-0.5 wt% of polymerization initiator, BPO. Drugs were incorporated into the polymer blend, followed by 15 seconds of ultrasonication to facilitate a homogeneous suspension. To initiate the crosslinking, the polymerization accelerator, DMT, was added in at a conetration of 0.1-0.5 wt % and thoroughly mixed, only after all other components had been added, resulting in the pre-polymer mixture. The mixture was drawn into a syringe and injected either into a sink for *in vitro* studies or subcutaneously into a rat for *in vivo* studies. Additionally, the mixture was pipetted into PDMS molds to create different shapes (cylinders, pipes, discs). The mixture underwent polymerization to form a solid depot within 3-10 minutes after the addition of the accelerator.

### Mechanical characterization

ISCD depots without or with TAF were prepared in cylindrical PDMS molds (6.8 mm diameter, 2.8mm height) for compression test. The dimensions of the depots were then measured using a caliper. The ISCD depots were mounted between plates of a mechanical tester (ADMET) and compressive force was applied to the samples at a rate of 1 mm/min. The compressive strain and stress on the samples were measured and the compressive moduli were obtained from the linear region (0.15-0.25 mm/mm strain) in the stress-strain curve. (n = 4)

### Rheological characterization

A rheometer (Discovery HR-3, TA Instruments) equipped with a parallel plate with a gap size of 1 mm and a diameter of 20 mm was used to characterize the viscosities of the samples and the injection force or to measure the cross-linking time.

To evaluate the injection force, PCLDMA without or with TAF (90 mg/mL) was pipetted onto the rheometer at room temperature and any excess solution was trimmed off with a spatula prior to measurements (n=3). The viscosity of the solution was measured while the shear rate was varied from 2 to 3700 s^−1^. The force required to inject 1 mL of ISCD solution in 30 seconds was measured indirectly from the viscosities using Hagen-Pouiselle equation^34^ as follows. Briefly, dynamic viscosities, μ, of ISCD solutions were measured and put into the equation below, where L is length of needle (15 mm); Q is volumetric flow rate (2 mL/min); D is inner diameter of syringe barrel (8.66 mm); and d is inner diameter of needle (0.337 mm).

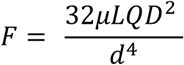

Briefly, dynamic viscosities, μ, of ISCD solutions were measured and put into the equation below, where L is length of needle (15 mm); Q is volumetric flow rate (2 mL/min); D is inner diameter of syringe barrel (8.66 mm); and d is inner diameter of needle (0.337 mm).

For crosslinking time measurements, PCLDMA was prepared with 0.1, 0.3, and 0.5 wt% BPO and mixed with 0.1, 0.3, and 0.5 wt% DMT, respectively. Immediately after mixing, the sample was pipetted onto the rheometer and the viscosity of the solution was monitored at a shear rate of 10 s^−1^ until the solutions were crosslinked (n=3). The time when the solutions started to increase in viscosity was recorded as the cross-linking time.

### Exothermic heat analysis

To quantify the exothermic heat generated during ISCD polymerization, 15 mg of PCLDMA was prepared with 0.1, 0.3, and 0.5 wt% BPO. The mixture was pipetted onto an aluminum pan and combined with 0.1, 0.3, and 0.5 wt% DMT, respectively. Immediately after mixing, the heat flow within the sample was measured for 20 minutes at room temperature. The total exothermic heat for the reaction was calculated by integrating the heat flow over time. For qualitative analysis, an infrared thermal imaging camera (Fluke Ti95 9Hz) was used to detect the temperature change in the depot during its polymerization.

### In vitro release

100 µl of Drug-loaded pre-polymer mixtures were injected into a sink medium. For TAC- and AMX-loaded ISCDs, we used 20% methanol in PBS as the sink medium due to low water solubility of these drugs. For other drugs, PBS was used as the sink. Sink containing ISCD depot was incubated in a shaker incubator at 37°C. To maintain sink conditions and prevent bacterial growth, the release medium was completely removed and replaced with fresh release medium every week. The release medium was collected at pre-determined time points to be analyzed directly or after lyophilization and reconstitution with an appropriate solvent for analysis.

*In vitro* cumulative release of TAF was analyzed using NMR. TAF release samples were stored in a −80°C freezer and lyophilized after completely frozen. The remaining powder was reconstituted with 500 µL 0.1 mg/mL maleic acid in deuterium oxide (D_2_O), as an internal standard, and loaded into NMR tubes (NORELL). NMR experiments were performed on an Agilent MR 400 MHz automated NMR system equipped with a 5mm AutoX One probe at room temperature. Sixty four scans were conducted for each 1H NMR spectrum recording. MestReNova was used for spectrum analysis. Baseline correction of the recorded spectra was performed manually in the software. The maleic acid alkene peak at approximately 6 ppm was integrated, whereas the ppm range of 8.1-8.3 (adenine protons) was integrated and normalized to the maleic acid peak integration area. A 3-point calibration curve was made in a range of 75-300 µg/mL of TAF in D_2_O with 0.1 mg/mL maleic acid. The concentration was plotted against the normalized area of integration. Linear regression was generated and used for all TAF sample quantification.

The *in vitro* cumulative release other drugs in this study was determined by HPLC (Agilent 1260 Infinity II) and the cumulative drug release was calculated. Sample analyses were performed on a ZORBAX 300SB-C18 column (Agilent, 3.0 x 150 mm, 3.5 µm) at 30°C, and all experiments were performed in triplicate. Samples were injected into the HPLC and chromatographic separation was achieved by gradient elution using different mobile phases and flow rates depending on the therapeutics, as described in **Table S1-S3**.

### Scanning electron microscopy (SEM)

Surface and microstructures of ISCD were imaged using SEM. First, pre-polymer mixtures with and without encapsulated drugs were prepared and polymerized in a PBS sink. At week 1 post-incubaiton, the implants were removed from the sink and dried overnight under low pressure. The lyophilized samples were subsequently mounted on an aluminum stub using carbon tape, and sputter coated with 6 nm of platinum (Leica EM ACE600). The coated samples were then imaged with Hitachi S-4700 FE-SEM.

### *In vitro* degradation analysis

To evaluate the degradation rate of different ISCDs, a weight loss assay was conducted at designated time intervals (Day1/2/4/7/14/28/30/42/60/80) at 37 °C. Depots were formed by injecting pre-polymer mixture into PBS (sink), followed by incubation in a shaker incubator at 37°C. Different samples were prepared for each time point. Sink was decanted at each pre-determined time point, and depots were subjected to freeze-drying for 24 hours to remove any residual sink solution. The weight loss (degradation) was quantified as a percentage according to the equation provided below.

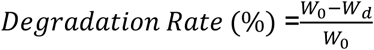

*W*_0_ is the original weight; *W*_*d*_ is the weight of the samples after degradation.

### Crosslinking density analysis

The degree of cross-linking of the depots was measured using an equilibrium swelling method and the Flory-Rehner equation, ^38^ described as follows, where *V*_*r*_ is the volume fraction of depot in swollen state; *V*_*s*_ is molar volume of the solvent (benzyl alcohol), 103.9 mL/mol; χ is the Flory-Huggins polymer-solvent interaction parameter.

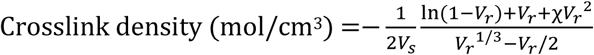

ISCD depots composed of PCLDMA with or without external polymer additives (PEG, PEGMMA, PEGDMA, PCL, or PDMS) with 0.3 wt% BPO/DMT and PCLDMA with 0.15 wt% BPO/DMT were prepared. The depots were incubated in benzyl alcohol for a week. The weight of each depot was measured before and after the incubation. The volume fraction of the depots in the swollen state was calculated from the increase in weight using the density of the solvent and polymer mixtures. The Flory-Huggins polymer-solvent interaction parameter was obtained from literature.^58^ The parameters were plugged into the Flory-Rehner equation to determine the cross-linking density of each depot.

### Animals

Animal experiments were conducted according to ethical guidelines approved by the Institutional Animal Care and Use Committee (IACUC) of Brigham and Women’s Hospital. Experiments were conducted in male Sprague-Dawley rats (6-8 weeks, Charles River Laboratories). Rats were maintained under pathogen-free conditions and randomly assigned to various experiment groups. The group size of animals in experiments was decided based on the minimum number of animals required to attain a statistical significance of *P*<0.05 among different test groups.

### Pharmacokinetics studies

PK studies were conducted with either unmodified ISCD or ISCD containing 25% PEGMMA. 0.3% BPO/DMT was used and the depots were loaded with TAF, NAL or TAC. The dosages used were 150 mg/kg for TAF and NAL, and 75 mg/kg for TAC. The drug-loaded pre-polymer mixture (500 µL) was administered subcutaneously with an 18 G needle over the shoulder of anesthetized rats. At pre-determined time points, blood was collected from the tail vein of the rats into EDTA-coated tubes.

For TAF and NAL analysis, blood samples were centrifuged at 1200g for 10 minutes at 4°C and the supernatant (plasma) was isolated. The Naltrexone/Nalbuphine Forensic ELISA Kit (Neogen) was used to quantify NAL plasma concentrations according to the manufacturer’s instructions. TAF and TFV concentrations in plasma samples were analyzed at the PPD Analytical Laboratory (PPD, Inc., Richmond, VA). A 25-μL matrix aliquot is fortified with a TFV-d6 and TAF-d5 internal standard working solution. TFV and TAF are isolated through a protein precipitation extraction. A portion of supernatant is evaporated under a nitrogen stream and the remaining residue is reconstituted. The final extract is analyzed via reversed phase chromatography and MS/MS detection using positive ion electrospray. A linear, 1/concentration² weighted, least-squares regression algorithm is used to quantitate samples.

To extract TAC from the blood samples, 100 µL of whole blood was combined with 100 µL of MeOH and 50 µL of 0.1M ZnSO4 in an Eppendorf tube, followed by vortexing. Subsequently, 1 mL of ethyl acetate was added to the mixture and vortexed again. The resulting mixture underwent centrifugation at 14000 rpm for 5 minutes at room temperature. The supernatant was collected and dried, and the measurement of tacrolimus levels was carried out using the Tacrolimus (FK506) ELISA kit (Abbexa, abx515779). The dried TAC sample was reconstituted with the sample diluent buffer provided in the ELISA kit, and further sample analysis was performed following the manufacturer’s instructions.

### PK analysis and human PK prediction

The PK analysis of ISCD formulations utilized both non-compartmental and compartmental methods. The in vivo release rate (mg/day) from ISCD formulations was determined employing the area function method^41^, a deconvolution technique that relies on the relationship between observed drug concentration following subcutaneous injection of ISCD and the area intervals under both subcutaneous and IV drug concentration-time curves. For compartment modeling of ISCD PK profiles in rats, the disposition kinetics of each drug were first established using rat IV PK data obtained experimentally (TFV, TAC) and from literature (NAL)^57–59^ with a two-compartment PK model. Subsequently, an appropriate absorption kinetic model was integrated to characterize the ISCD PK profiles. These absorption models describe the in vivo release rate profiles of ISCD, characterized by multiple first-order rates occurring in a sequential manner.

For human PK prediction for NAL and TAC following subcutaneous injection of ISCD, convolution analysis was employed, integrating the human disposition kinetic function and the release function of the ISCD formulation. The human disposition function was derived from human IV PK data for TAC^48^ and NAL^47^ obtained from the literature. The in vivo release functions estimated from the rat PCLDMA ISCD were used after applying simple allometry to release kinetic parameters based on average body size between rats and humans, with a typical allometry exponent of −0.25 for rate constants^60, 61^.

### Elemental CHN analysis

Residual TAF content within the depots was quantified using elemental analysis (CHN) performed at Midwest Microlab (Indianapolis, IN). Given that the polymer matrix used in the depots lacks nitrogen, the measured nitrogen content can be directly correlated with the remaining amount of TAF. The mass percentages of carbon, hydrogen, and nitrogen in the sample were analyzed, and the residual drug mass within the depots was back-calculated from these values.

### *In vivo* degradation

To assess *in vivo* degradation, weight loss of explanted ISCDs was measured. Rats received subcutaneous injections of 500 µL unmodified ISCDs or ISCDs containing 25% PEGMMA (both with 0.3% BPO/DMT but no drugs) (n=3). After 7 months, the rats were euthanized, and the explanted depots were cleared of surrounding tissue. The explants were rinsed with DI water and freeze-dried for 24 hours to remove residual solvents. Weight loss was then calculated as a percentage of degradation using the equation provided below.

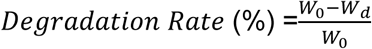

### *In vivo* biocompatibility study

To investigate the inflammatory response caused by the implanted material, rats received injections of either unmodified ISCD or ISCD containing 25% PEGMMA, both formulated with 0.3% BPO/DMT but without any drugs. The depots were explanted at 1 week, 4 weeks, and 7 months post-injection. To assess tissue response over time, histological and immunohistochemical analyses were performed on cryosections of the explanted samples. After explantation, samples were fixed in 4% paraformaldehyde for 4 hours, followed by overnight incubation in 30% sucrose at 4 °C. Samples were then embedded in optimal cutting temperature compound (OCT) and flash frozen in liquid nitrogen. Frozen samples were then sectioned using a Leica Biosystems CM1950 Cryostat. 15-μm cryosections were obtained and mounted in positively charged slides. The slides were then processed for hematoxylin and eosin staining (Sigma) according to instructions from the manufacturer. The stained samples were preserved with DPX mountant medium (Sigma). Immunohistofluorescent staining was performed on mounted cryosections. Anti-CD3 (ab16669) and anti-CD68 (ab125212) (Abcam) were used as primary antibodies, and an Alexa Fluor 594-conjugated secondary antibody (Invitrogen) was used for detection. All sections were counterstained with DAPI (Invitrogen), and visualized on an Leica DMi8 widefield microscope.

### Depot retrievability

We subcutaneously administered 500 µL ISCD composed of PCLDMA, 90 mg/mL TAF and 0.3 % BPO/DMT with an 18 G needle to rats and retrieved the depots two weeks later. For the retrieval, the rats were anesthetized and their back was shaved to visualize the location of the depot. Under sterile conditions, a small cutaneous incision was made adjacent to the depot for atraumatic removal using forceps, followed by closure with sterile sutures. Blood (300-500 µL) was collected one hour before retrieval and at day 1, 3, 7, 10, 14, and 21 post-retrieval to monitor plasma TFV levels.

### Statistical information

All values are presented as mean ± standard deviation. Two-tailed Student’s t-test was used to compare two experimental groups, and one-way ANOVA with Tukey’s post hoc analysis was used to compare more than two groups using GraphPad (Software Inc., CA, USA), with P values defined as *< 0.05, **< 0.01, ***< 0.001, and ****< 0.0001. One-way ANOVA with Tukey’s post hoc analysis was used to determine the statistical significance of differences in swelling and cross-linking density of ISCD, with the mean of each group compared to the mean of the PCLDMA with 0.3% w/w BPO/DMT (control) group. To compare the *in vivo* plasma TFV level of unmodified ISCD and ISCD containing 25% PEGMMA, we used a two-way ANOVA analysis with time and ISCD formulation as the two variables. One-way ANOVA with Tukey’s post hoc analysis was used to determine the statistical significance of differences in residual TAF from explanted depots, with the mean of each group compared to the mean of the unmodified ISCD depot.

## Supporting information

ISCD_Supplementary Information

## Acknolwedgements

We acknowledge funding support from National Institute of Health Grant R21DA057701 (to NJ) and Department of Anesthesiology, Perioperative, and Pain Medicine at the Brigham and Women’s Hospital (to NJ and JMK).

## Author contribution

Conceptualization: SL, SZ, JMK, NJ

Methodology: SL, SZ, WJ, BB, SW, NJ

Investigation: SL, SZ, WJ, XC, LZ, JJ, EA, HBM, SB, KS, KC, JNL, CS, JG, DL, CJ, AG

Supervision: SW, JMK, NJ

Writing—original draft: SL, WJ, SW, NJ

Writing—review & editing: SL, JMK, NJ

## Supplementary Information

Supplementary Information: should be combined and supplied as a separate file, preferably in PDF format.

