## Supplementary material for "Ultra-Long-Term Delivery of Hydrophilic Drugs Using Injectable *In Situ* Cross-Linked Depots": ISCD_Supplementary Information

**This PDF file includes:**

**Supplementary Figure 1 to 13**

**Supplementary Tables 1 to 6**

### SUPPLEMENTARY FIGURES

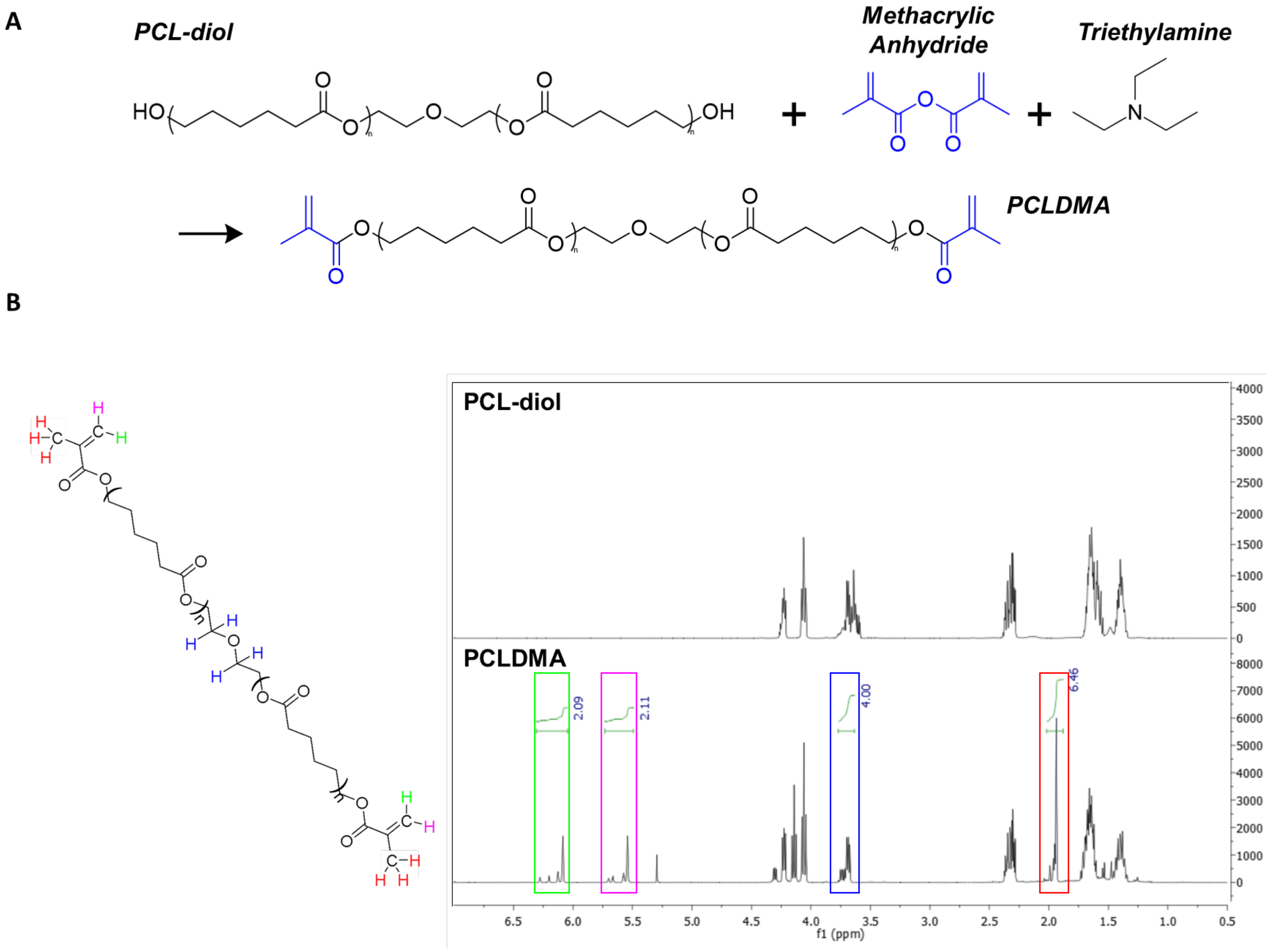


**Figure S1. Synthesis and characterization of PCLDMA. A.** Schematic showing methacrylation of hydroxyl-functionalized PCL to yield PCLDMA. **B.** $^{1}$H-NMR spetrcum of of PCLDMA. The chemically equivalent protons are labeled with the same color and their NMR signals are marked with the same colored box for ease of understanding. The NMR integration of each signal corresponds to the expected ratio of each type of hydrogen in the PCLDMA molecule.

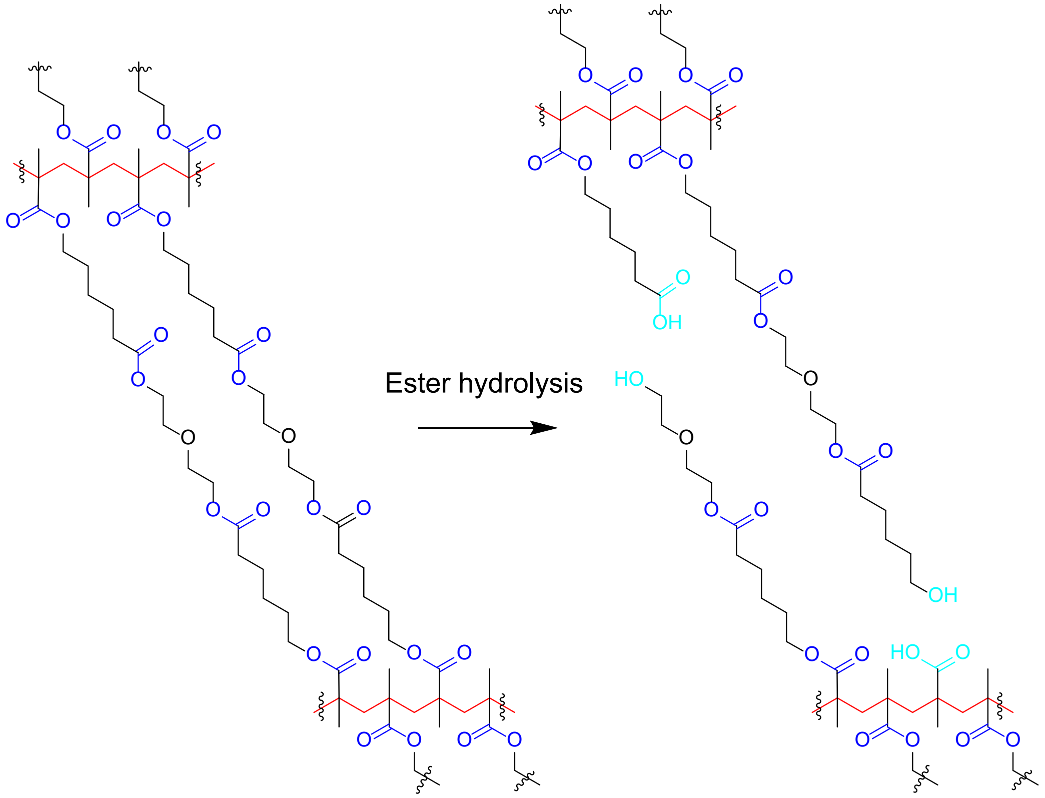


**Figure S2.** Schematic illustrating hydrolysis of polycaprolactone domains and subsequent collapse of ISCD. The ISCD consists of a cross-linked network of polymethacrylate chains (red) connected by ester bonds (dark blue). Hydrolysis of these ester bonds (light blue) disconnects the polymethacrylate chains, enabling the depot to degrade.

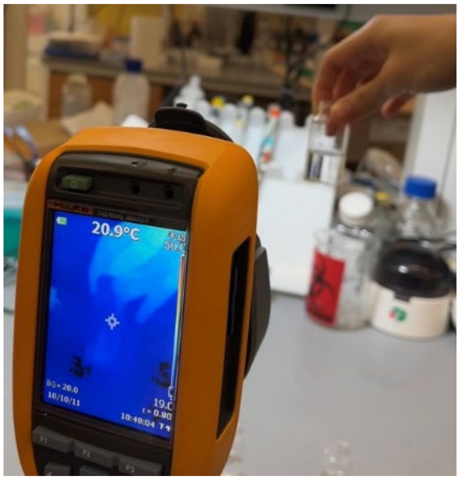


**Figure S3.** Infrared thermal imaging confirms that there is no noticeable heat generated during ISCD polymerization.

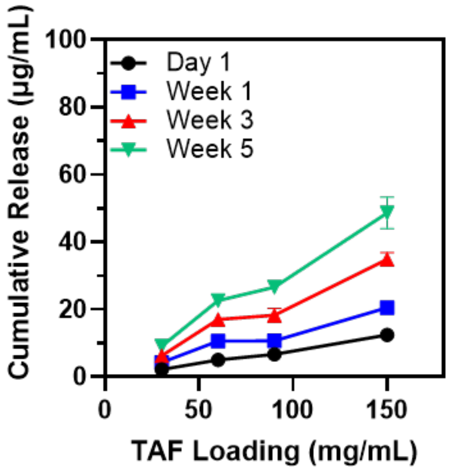


**Figure S4.** *In vitro* cumulative release of TAF from ISCD loaded with different concentrations of TAF and incubated in PBS (37°C). Data are presented as mean ± standard deviation (n=3, experiments performed at least twice).

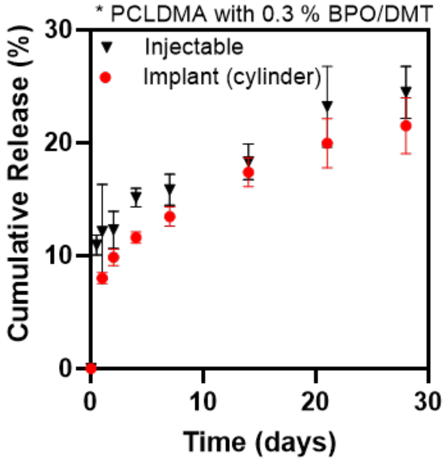


**Figure S5.** *In vitro* cumulative release of TAF from ISCD depots formed by injecting pre-polymer mixture into PBS (37°C) compared with TAF release from pre-formed implants with cylindrical shape. Data are presented as mean ± standard deviation (n=3, experiments performed at least twice).

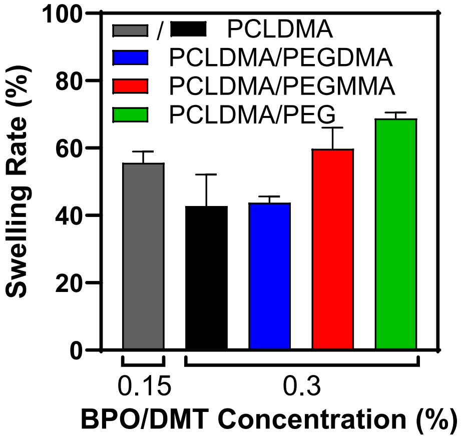


**Figure S6.** Swelling rate of different ISCD formulations studied in benzyl alcohol aftera week. Data are presented as mean ± standard deviation (n=3, experiments performed at least twice).

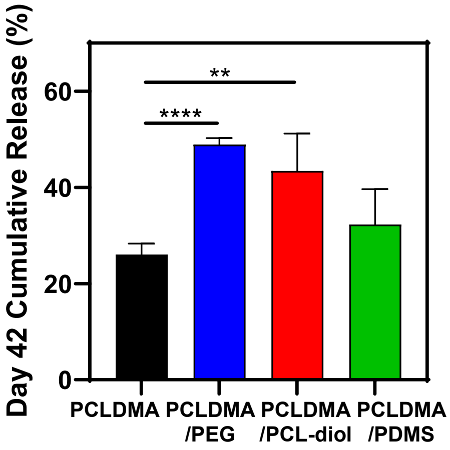


**Figure S7.** Cumulative release of TAF at day 42 post-incubation of different ISCD formulations in PBS (37$^{o}$C). (**P<0.01, ****P<0.0001). Data are presented as mean ± standard deviation (n=3). The P-value was determined using one-way ANOVA with Tukey's post hoc analysis.

.

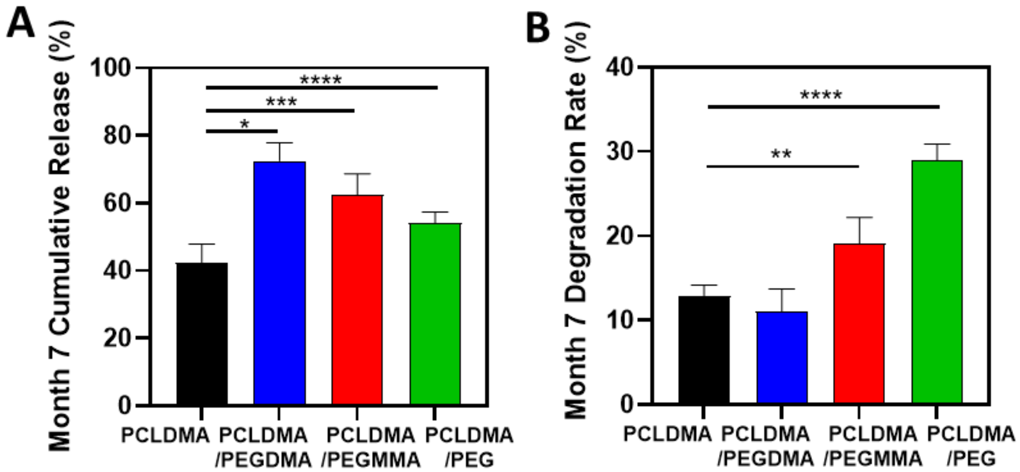


**Figure S8. Impact of incorporating external polymer additives with varying degrees of methacrylation into ISCD on TAF release and depot degradation. A.** Month 7 cumulative release, and B. month 7 degradation rate of unmodified ISCD and ISCD containing 25 wt% of PEGs with different degree of methacrylation, when incubated in PBS (37°C). (*$P<0.05$, **$P<0.01$, ***$P<0.001$, ****$P<0.0001$). Data are presented as mean ± standard deviation (n=3). The *P*-value was determined using one-way ANOVA with Tukey's post hoc analysis.

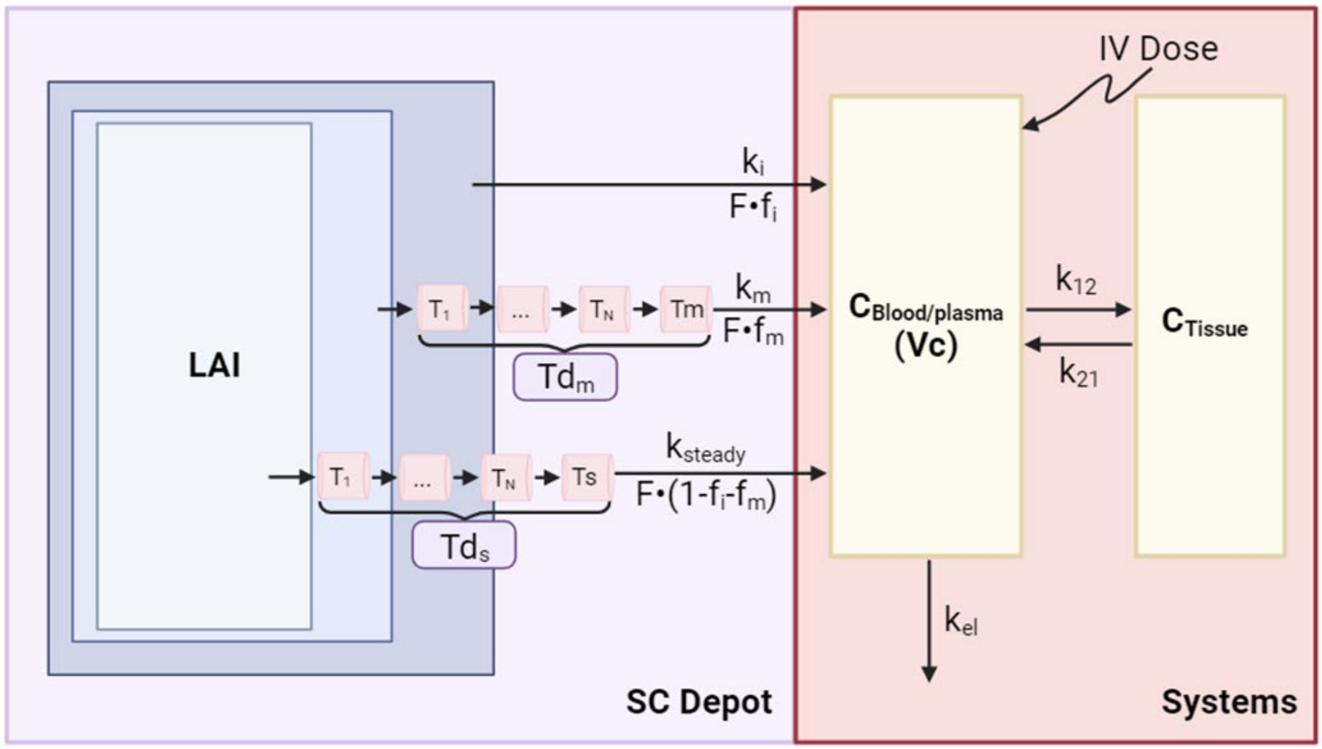


**Figure S9. The scheme of the pharmacokinetic (PK) model for subcutaneous injection of ISCD.** The disposition kinetics (referred to as Systems) of analytes is characterized by two compartments ($C_{Blood/Plasma}$ and $C_{Tissue}$), with first-order rate constants for elimination ($k_{el}$), distribution ($k_{12}$), and redistribution ($k_{21}$), and Vc for the central volume of distribution. At the SC implant site (referred to as SC Depot), the release/absorption model assumes three sequential release phases, delineated by first-order release rate constants ($k_i$, $k_m$, $k_s$). Initially, a fraction ($f_i$) of the ISCD implant is released ($k_i$), leading to the maximum concentration in the central compartment. Subsequently, drug release continues with an intermediate phase ($k_m$) for a fraction of the total released drug mass ($f_m$), followed by a sustained release phase ($k_s$) for the remaining drug amount (1- $f_i$- $f_m$). The time delays associated with the intermediate (Tdm) and sustained-release (Tds) phase are characterized by a gamma distribution function with shape (N) and rate parameter (Td). F represents the bioavailability of ISCD implants.

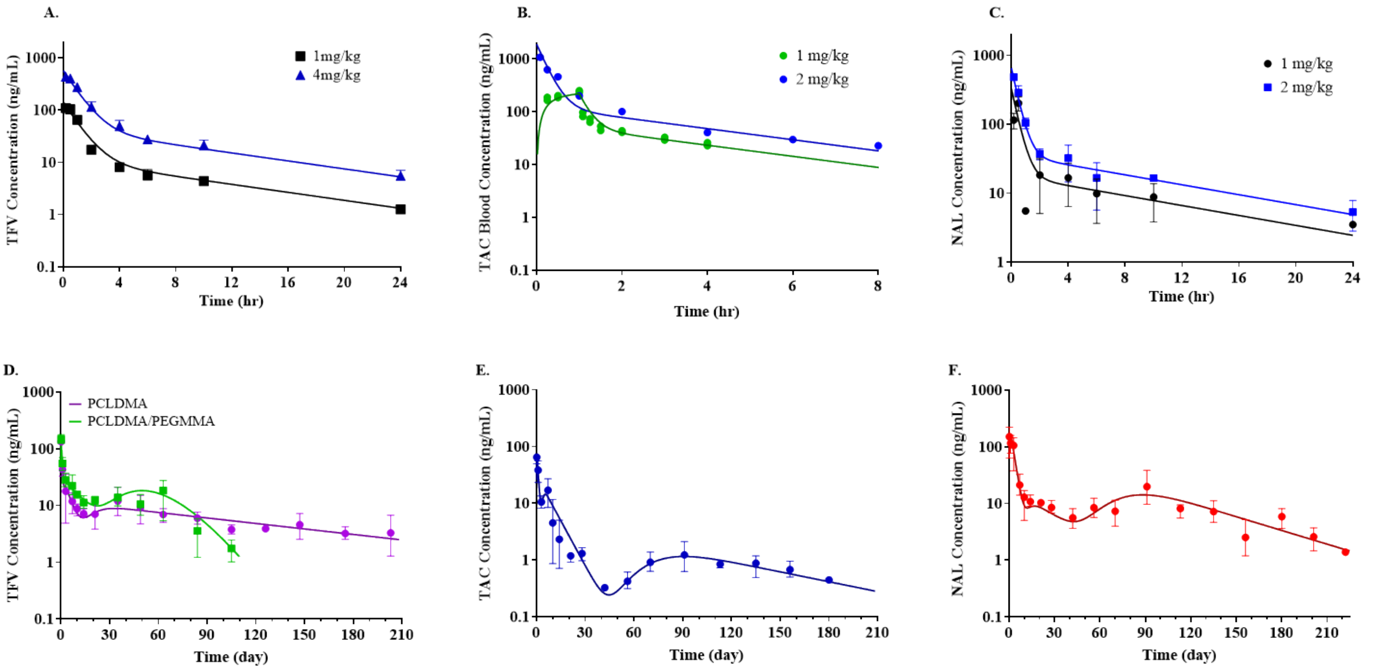


**Figure S10.** Time profiles of the blood/plasma concentration in rats after a single IV dose of **A.** TFV (1 & 4 mg/kg) **B.** TAC (1 & 2 mg/kg), and **C.** NAL (1 & 2 mg/kg). Time profiles of the blood/plasma concentration in rats after subcutaneous injection of ISCD containing **D.** TFV, **E.** TAC, and **F.** NAL. Data presented as symbols reflect the mean ± standard deviation of three technical repeats (n=3). Lines represent the model-predicted drug concentrations.

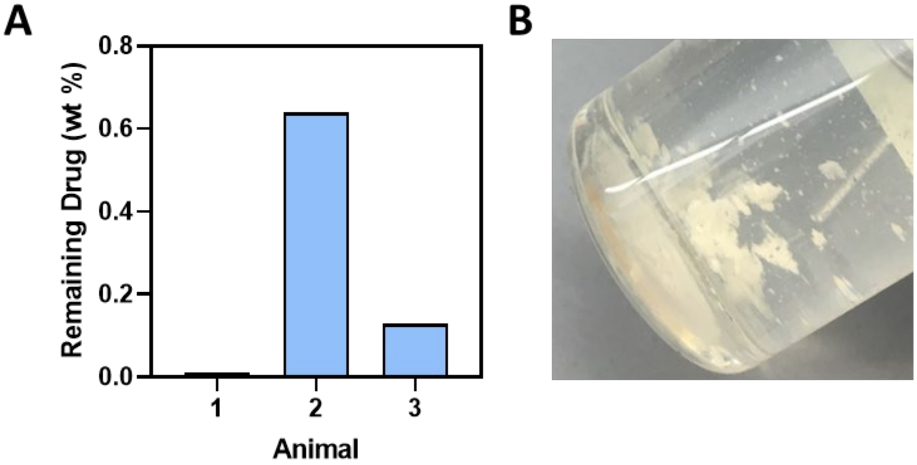


**Figure S11 A.** Pecentage of the initial amount TAF amount remaining in the explanted ISFI following the 2-month *in vivo* study in rats. B. ISFI explanted after the 2-month *in vivo* study is a fragmented solid. Data are presented as individual values for each animal.

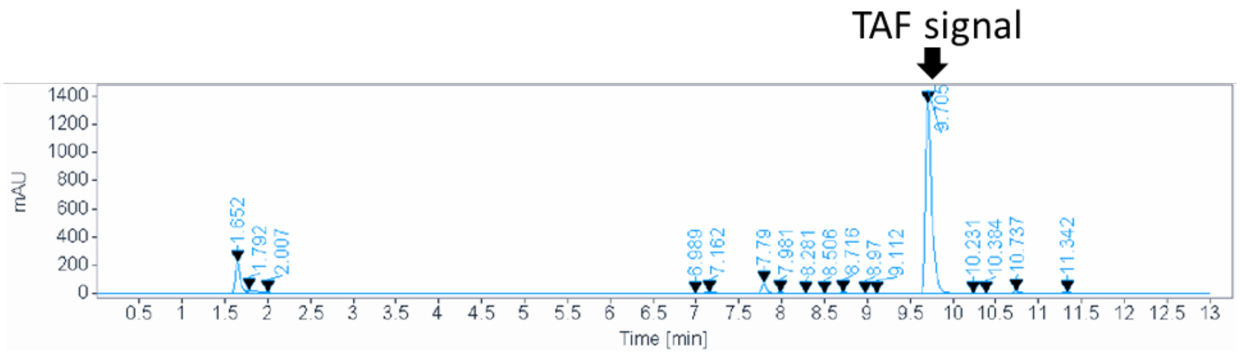


**Figure S12.** HPLC chromatograph of TAF-loaded ISCD explanted after a 7-month *in vivo* study.

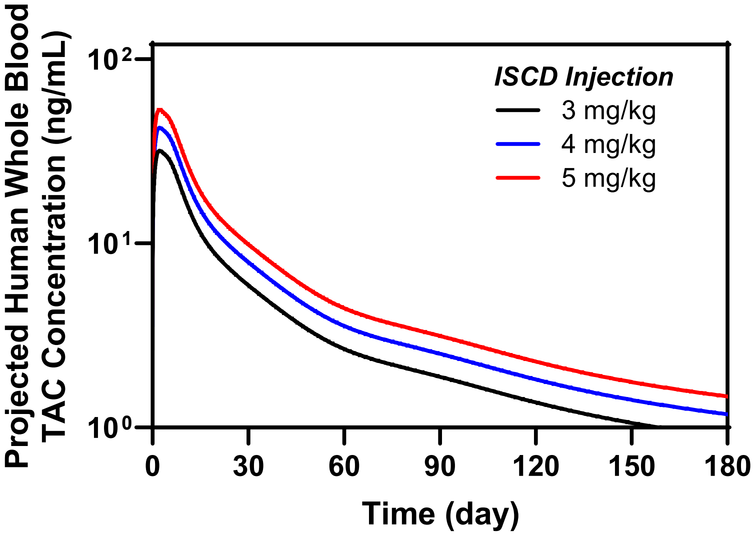


**Figure S13.** Convolution analysis-based prediction of human PK of a single subcutaneous dose of TAC-loaded ISCD (at different dosages) upto 6 months.

### SUPPLEMENTARY TABLES

**Table S1. Initial burst release against water solubility of therapeutics**

| Drug | Solubility (mg/ml) | Cumulative release after 24 h (%) |
|---|---|---|
| FTC | 112 | 18.51 |
| NAL | 100 | 22.02 |
| LAM | 70 | 14.5 |
| VAN | 50 | 14.98 |
| TAF | 5.63 | 12.14 |
| AMX | 3 | 0.79 |
| ABC | 1.21 | 6.04 |
| TAC | 0.004 | 1.34 |

**Table S2.** The estimated PK parameters of disposition and release kinetics were obtained from the concentration-time profiles following IV administration and SC implants of ISCDs in rats.

| **Parameters** | **Definition** | **PCLDMA +TAF** | **PCLDMA/ PEGMMA +TAF** | **PCLDMA +TAC** | **PCLDMA +NAL** |
|---|---|---|---|---|---|
| *Disposition Kinetics* | | | | | |
| **$V_c$ (mL)** | Central compartment volume | 1266 (9)[a] | | 306 (17) | 923 (20) |
| **$k_{el}$ (1/hr)** | Elimination rate constant | 0.558 (8) | | 2.06 (16) | 0.925 (17) |
| **$k_{12}$ (1/hr)** | Transfer rate constant from central to peripheral compartment | 0.372 (15) | | 2.08 (21) | 1.22 (19) |
| **$k_{21}$ (1/hr)** | Transfer rate constant from peripheral to central compartment | 0.158 (12) | | 0.517 (21) | 0.203 (17) |
| **$t_{1/2}$ (hr)** | Terminal elimination half-life | 7.88 | | 2.87 | 8.35 |
| **CL (L/hr)** | Total systemic clearance | 707 | | 631 | 853 |
| *Release Kinetics* | | | | | |
| **$k_i$ (1/day)** | Initial release rate constant | 2.47 (27) | 2.52 (25) | 0.962 (43) | 0.372 (13) |
| **$f_i$** | Fraction of the released drug mass associated with ki | 0.058 (18) | 0.079 (17) | 0.231 (26) | 0.274 (11) |
| **$k_m$ (1/day)** | Intermediate release rate constant | 0.178 (34) | 0.0643 (27) | 0.122 (8) | 0.0346 (64) |
| **$f_m$** | Fraction of the released drug mass associated with km | 0.092 (18) | 0.315 (16) | 0.413 (15) | 0.154 (38) |
| **Tdm (day)** | Mean transit time for drug release associated with km | 1.5[b] | 1[b] | 3.7 (29) | 11.8 (24) |
| **$N_m$** | Number of transit compartment for drug release associated with km | 5[b] | 5[b] | 10[b] | 10[b] |
| **$k_s$ (1/day)** | Sustained release rate constant | 0.00744 (16) | 0.128 | 0.0134 (28) | 0.0181 (16) |
| **Tds (day)** | Mean transit time for drug release associated with ks | 18.1 (23) | 50.8[b] | 68.3 (7) | 70.1 (9) |
| **$N_s$** | Number of transit compartment for drug release associated with ks | 5[b] | 6[b] | 15[b] | 15[b] |
| **$F_{total}$** | Projected total bioavailability of ISCD | 1[b] | 0.86 (7) | 0.38 (7) | 1[b] |
| **$F_{tlast}$** | Fraction of the total drug mass until the last observed concentration | 0.80 | 0.86 | 0.35 | 0.97 |

[a] Coefficient of Variability (CV)%; [b] Fixed parameters

**Table S3.** The disposition PK parameter values obtained from the concentration-time profiles of TAC (0.075 mg/kg/day) and NAL (1 mg) following oral or IV administration in humans.

| **Parameter (unit)** | **TAC** | **NAL** |
|---|---|---|
| **CL (L/hr)** | 3.3 | 248 |
| **$V_{ss}$ (L)** | 101 | 563 |
| **$k_{12}$ (1/hr)** | 0.276 | 3.1 |
| **$k_{21}$ (1/hr)** | 0.131 | 1.84 |
| **$t_{1/2}$ (hr)** | 24.8 | 1.83 |

**Table S4. HPLC method details for different therapeutics**

| Therapeutic compound | Flow rate (mL/min) | Retention time (min) | Mobile phase | Gradient elution | Detection wavelength (nm) | Injection volume (μL) |
|---|---|---|---|---|---|---|
| TAC | 1 | 19 | methanol/DI water (70:30 v/v) | Table S5 | 210 | 20 |
| LAM, ABC, NAL | 0.45 | 13 | 10mM ammonium formate buffer/ACN (95:5 v/v) | Table S6 | 220-NAL<br>259-ABC<br>271-LAM | 5 |
| AMX | 1 | 8 | 25 mM phosphate buffer/ACN (95:5 v/v) | None | 240 | 20 |
| VAN | 1 | 8 | 20mM ammounium acetate buffer/methanol (88:12 v/v) | None | 240 | 20 |

**Table S5. Gradient program of the mobile phase for HPLC analysis of TAC**

| Time (min) | Mobile Phase | |
|---|---|---|
| | Methanol (%) | Water (%) |
| 3 | 70 | 30 |
| 15 | 90 | 10 |
| 16 | 90 | 10 |
| 16.1 | 70 | 30 |
| 19 | 70 | 30 |

**Table S6. Gradient program of the mobile phase for HPLC analysis of LAM, ABA, and NAL**

| Time (min) | Mobile Phase | |
|---|---|---|
| | Methanol (%) | Water (%) |
| 2 | 95 | 5 |
| 3 | 95 | 5 |
| 7 | 40 | 60 |
| 8 | 40 | 60 |
| 8.1 | 95 | 5 |
| 3 | 95 | 5 |
